# High-resolution cryo-EM of ribosomes in mammalian neurons

**DOI:** 10.64898/2026.11.25.769232

**Authors:** Stephen Diggs, Johannes Elferich, You Zhou, Soojin Lee, Ximena Zottig, Zahra Seraj, Dilara Halim, Fen-Biao Gao, Nikolaus Grigorieff, Andrei A. Korostelev

**Affiliations:** RNA Therapeutics Institute, UMass Chan Medical School, 368 Plantation Street, Worcester, MA 01605, USA; Howard Hughes Medical Institute, UMass Chan Medical School, 368 Plantation Street, Worcester, MA 01605, USA

**Author notes:** School of Biopharmaceutical and Medical Sciences, Sungshin Women’s University, Seoul, 01133, Korea; Basic Science Institute, Sungshin Women’s University, Seoul, 01133, Korea (Soojin Lee).

**Keywords:** Decoding center, DeCo-LACE, iPSC, PTC, Neuronal translation, 2DTM, uL16

## Abstract

Brain function depends on protein synthesis (translation), which controls neurodevelopment, synaptic plasticity, and memory formation. To understand the molecular mechanisms of translational control, the functional states of neuronal ribosomes must be determined. We visualized the translational apparatus within induced pluripotent stem cells (iPSC)-derived human neurons and in human and mouse neuron-derived lysates, using cryogenic electron microscopy (cryo-EM) with high-resolution 2D template matching (2DTM) and large-area imaging (DeCo-LACE). Cryo-EM maps at up to 2.6 Å resolution reveal structural details of numerous functional states of ribosomes. Initiating 40S subunits abundantly feature initiation factor eIF1S (also known as 5MP) indicating a tight control of translation initiation. Ribosomes with elongation factors comprise ∼90% of 80S particles, suggesting a high prevalence of translating ribosomes. Most elongating ribosomes sample pre-decoding states with aminoacyl-tRNA•eEF1A•GTP and a well-resolved decoding center prior to aminoacyl-tRNA acceptance. On the large ribosomal subunit, a conserved loop of uL16 stabilizes the CCA end of peptidyl-tRNA, suggesting a ribosomal protein contribution into RNA-catalyzed peptidyl transfer. The composition of peripheral ribosomal proteins, including the P-stalk proteins proposed to be substoichiometric in neurons, is homogeneous throughout ribosome states. The compositional heterogeneity of ribosomes is therefore negligible or only occurs under specific cellular conditions. Our work provides a comprehensive visualization of ribosome states within neurons and paves the way for probing translation regulation and dysregulation in neurological diseases using cryo-EM.

**Significance:** Neurons rely on tightly controlled protein synthesis to develop, learn, and allow formation of memories, yet how their ribosomes, the cell’s protein factories, are regulated remains largely unknown. Using cryo-electron microscopy to image ribosomes directly in human neurons and neuronal extracts, we show that most ribosomes are actively elongating, and that initiating ribosomes are almost fully occupied by eIF1S, a protein factor that enforces start-site stringency, pointing to highly efficient and accurate neuronal translation. We find no evidence for previously suggested neuron-specific “specialized” ribosomes lacking P-stalk proteins, emphasizing mostly canonical translation regulation. Our high-resolution structures also reveal new details of the decoding mechanism. This work provides a molecular framework for studying the role of translation dysregulation in neurological disease.

## Introduction

Protein synthesis (translation) governs all aspects of brain development and function, including synaptic plasticity and memory formation (1, 2). Early neurobiology studies, for example, established that *de ovo* protein synthesis is essential for long-term memory formation. Consequently, when translating ribosomes are inhibited, the consolidation of new memories is blocked (1, 3, 4). Accordingly, dysregulation of translation in neurons contributes to numerous neurodevelopmental and neurodegenerative diseases (5–8). The majority of ribosomes in neurons reside in the soma, where they produce most cellular proteins, and a fraction of the ribosomes are transported to axons and dendrites, where translation underlies axonal guidance, dendritic growth, and synaptic plasticity (1, 2, 9). Translation is highly regulated to control protein output. To achieve this, ribosomes interact with a large array of protein factors and undergo conformational and compositional rearrangements at all stages of translation. For example, elongating ribosomes experience intersubunit rotation as they exchange elongation factors while moving along an mRNA (10–13). Alternatively, unused (hibernating) ribosomes that accumulate during stress are devoid of mRNA and bind different sets of proteins (14, 15). Distinguishing among ribosomal conformational states and occupancies of factors, tRNAs, and mRNA (collectively, “ligands”) can therefore inform on the extent of protein synthesis and roles of regulatory factors under different conditions.

To understand neuronal translation regulation, it is essential to resolve detailed structures of ribosomes in neurons. Recent progress in cryogenic electron microscopy (cryo-EM) has enabled atomic-resolution structure determination from *in vitro* reconstituted complexes, and new developments, most notably in cryogenic electron tomography (cryo-ET), continue to improve structure resolution within cellular contexts (10, 16–19). While *in situ* cryo-ET allows the distinction of individual subunits and ribosome particles, the resolution of tomograms is usually limited to ∼10 Å, which does not allow unambiguous identification of ligands and detailed structural differences in ribosomes (20, 21). In *ex vivo* cryo-EM studies, such as analyses of polysomes or ribosomes isolated from cell cultures or tissues (22–25), near-atomic resolutions can be achieved, but ribosome state distributions and translation factor occupancy may change due to the disruption of the cellular environment and purification procedures (14). Furthermore, high-resolution cryo-EM studies performed primarily on immortalized cell lines, or cells that have not been terminally differentiated, have left unaddressed how terminally differentiated cells like human neurons control translation. The compositions of ribosome complexes and functional state distributions in functional neurons therefore remain unknown.

Here, we describe high-resolution cryo-EM analyses of mammalian neuronal translation, using *in situ* cryo-EM and *in extracto* cryo-EM of the translational machinery in the presence of cytoplasmic components (14, 26). We collected cryo-EM data from several sources: cryo-FIB milled human iPSC-derived motor neurons (hiPSC-MN), lysates obtained from hiPSC-MN, and lysates obtained from mouse primary cortical neurons (mouse PCN). High-resolution 2D template matching (2DTM) enabled efficient detection of ribosome particles in crowded cellular environments (27–29), and defocus-corrected large-area cryo-EM (DeCo-LACE) allowed for accurate mapping of ribosomal functional states directly in hiPSC-MN lamella (30). Extensive sub-classification yielded dozens of 40S, 60S, and 80S reconstructions in distinct functional states (31, 32). Elongating ribosomes represent the largest population, suggesting at least some neuron types achieve highly efficient translation (33, 34). High-resolution analyses reveal a uL16 protein loop as the conserved feature at the peptidyl transferase center and interactions of the P stalk with elongation factor eEF1A, delineating important functions of ribosomal proteins in protein synthesis and mRNA decoding.

## Results and Discussion

### Predominance of elongating ribosomes in hiPSC-MN lamellae

To investigate ribosomes in neurons, we prepared FIB-milled lamellae from 30-day-old terminally differentiated hiPSC-MN. To be selected for FIB milling and cryo-EM data collection, intact neurons first had to pass a cell-viability assay (*SI Appendix*, Fig. S1 *A-F*). Thinned lamellae were imaged using the DeCo-LACE method, yielding high-resolution micrographs of large cell areas (Fig. 1*A* and *SI Appendix*, Fig. S1*G*) (30). To enable particle detection in these images, high-resolution 2DTM was performed using the human 60S ribosomal subunit as a template (*SI Appendix*, Fig. S2*A*; (35), PDB: 6QZP). Lamellae containing a sufficiently large portion of cytoplasm yielded several hundred matches per lamella (Figs. 1*A* and *SI Appendix*, Fig. S1).

**Figure 1.**
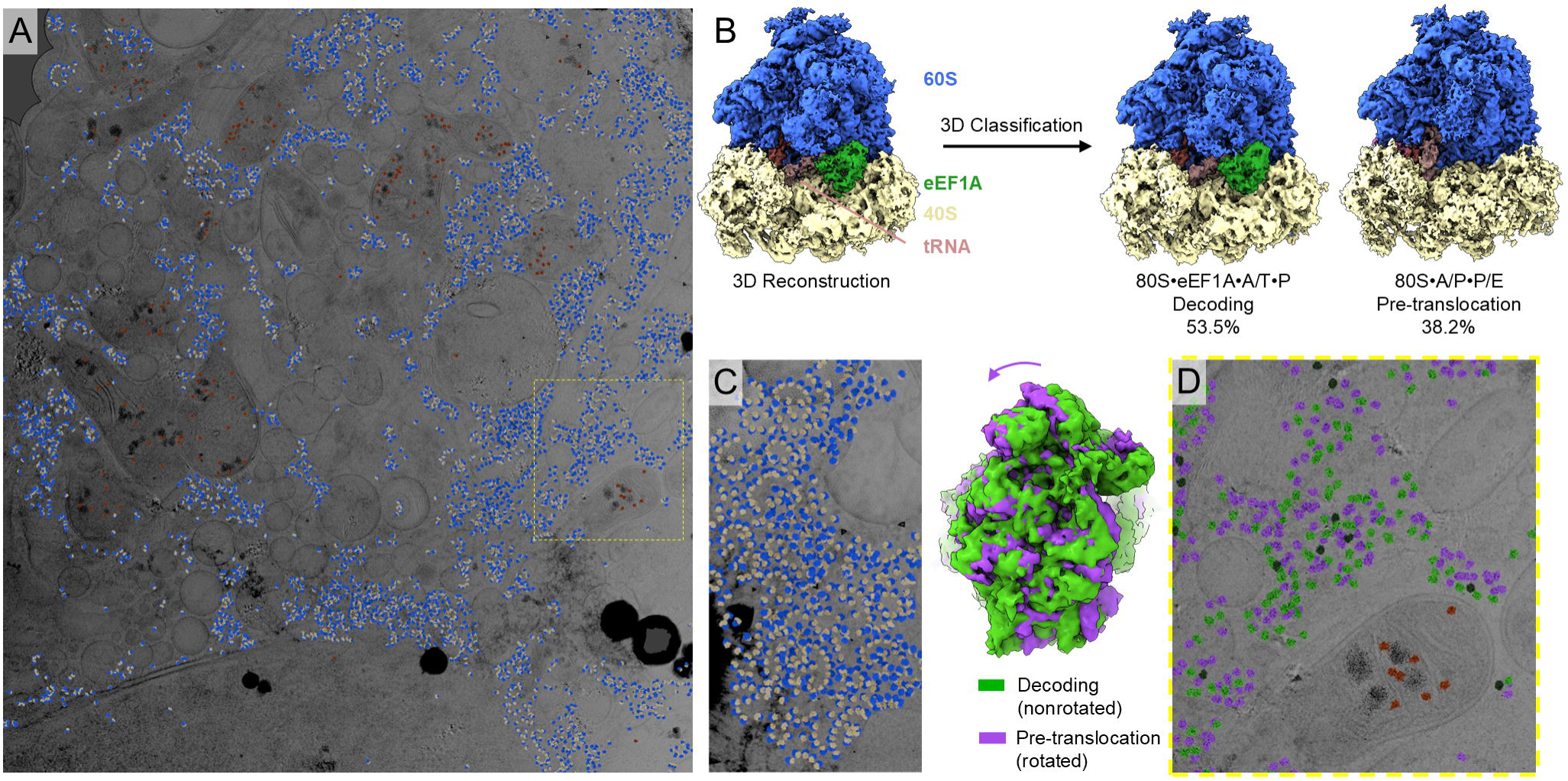
*In situ* cryo-EM of hiPSC-MN via 2DTM. (A) A fragment of DeCo-LACE montage overlaid with cytoplasmic ribosomes in blue and yellow, and mitochondrial ribosomes in orange (see *SI Appendix*, Fig. 1G for the complete montage). (B) 3D reconstruction after 2DTM (left) and the two predominant classes resulting from 3D classification representing the eEF1A-bound “decoding” (with A/T and E-tRNAs) and pre-translocation rotated (with hybrid-state A/P and P/E tRNAs) states. (C) Potential polysome organization within lamella. (D) Mapping of ribosome particle class assignment onto micrographs showing the cellular localization of functional ribosome states. Decoding and pre-translocation states are in green and purple, respectively, lower-resolution particles are in dark green.

Ribosomes often appeared in clusters (Fig. 1C), in agreement with previous reports that the soma mostly contains polysomes (9). Certain regions devoid of cytoplasmic ribosomes could be explained by their compartmental identity, such as the interior of the nucleus (Fig. 1*A*, bottom middle). Furthermore, mitochondria could be differentiated from cytoplasmic compartments by matching mitochondrial ribosomes, emphasizing the selectivity of 2DTM (Fig. 1*A*).

The initial cryo-EM map, calculated from ∼80,000 particles, showed strong density for the 80S ribosome, elongation factor 1A (eEF1A), A/T tRNA, and P-site tRNA (Fig. 1*B*, left), indicating that elongating ribosomes form a predominant population. Indeed, extensive classification of the dataset (36) resolved two elongation states. The most abundant is the “decoding” state featuring eEF1A delivering A/T tRNA to the A site, and P-site tRNA, accounting for 53.5% of particles (Fig. 1*B*, middle left). The “pre-translocation” state (38.2%) features a rotated 80S conformation, in which the 40S subunit is rotated relative to the 60S subunit, with two hybrid-state tRNAs in the A/P and P/E positions (the first letter denotes the position on the 40S subunit and the second on the 60S subunit; Fig. 1*B*, right). The remaining lower resolution states contained features resembling eEF1A-like GTPase and P-site tRNA, likely representing heterogeneous mixtures of elongating and/or other ribosome states. All reconstructions contain strong density for ErbB3-binding protein 1 (EBP1), which was proposed to associate with ribosomes under stress, indicating that EBP1 is also bound to the majority of elongating ribosomes in neurons, echoing recent findings for immortalized cells (14, 37–39) (*SI Appendix*, Fig. S2 *B-E*). Elongating ribosomes also feature density consistent with the C-terminal tail of hibernation factor SERBP1 stretching from eS10 at the 40S beak through uS10 to RACK1, where it is bound via the FPxL motif, similar to that in HeLa cells (38). Here, SERPB1 likely protects the elongating ribosomes from stalling-induced stress-responses and ribosome degradation (40). In summary, the elongating ribosomes represent more than 90% of neuronal ribosomes (Fig. 1*B* and *SI Appendix*, Fig. S2 *F-G*), consistent with decoding and pre-translocation being the rate-limiting steps, as the ribosome pauses to bind a cognate tRNA or elongation factor eEF2 (34, 41).

Recent studies showed that immortalized cell cultures contain substantial populations (>25%) of inactive ribosomes without mRNA, also known as hibernating ribosomes, as well as individual 60S subunits disengaged from translation (14, 38). By contrast, no hibernating 80S or isolated 60S subunits were detected in neuron lamellae, indicating that these inactive states, if present, account for a much lower population of ribosomes than in other cell types. To gain deeper insights into neuronal ribosomes and separate states that could not be resolved in the lamella dataset, we employed *in extracto* cryo-EM (14), allowing us to obtain near-atomic resolution of numerous additional states from larger datasets, including low-abundance states.

### In extracto cryo-EM of hiPSC-MN and mouse PCN

To visualize neuronal ribosomes at higher resolution, we performed *in extracto* cryo-EM on 30-day-old hiPSC-MN. This approach allows fast collection of large datasets from quickly lysed cells, thus bypassing lamellae preparation, and results in high-resolution reconstructions of the ribosome in the presence of cellular components, yielding cell-like representation of functional states (14). In parallel, immunofluorescence assays confirmed that neurons did not undergo stress or apoptosis (Fig. 2*A* and *SI Appendix*, Fig. S3). Upon cell lysis, the extracts were vitrified in liquid ethane, and data were collected as for single-particle cryo-EM and processed using 2DTM (14) (see Supplementary Methods and *SI Appendix*, Fig. S4).

**Figure 2.**
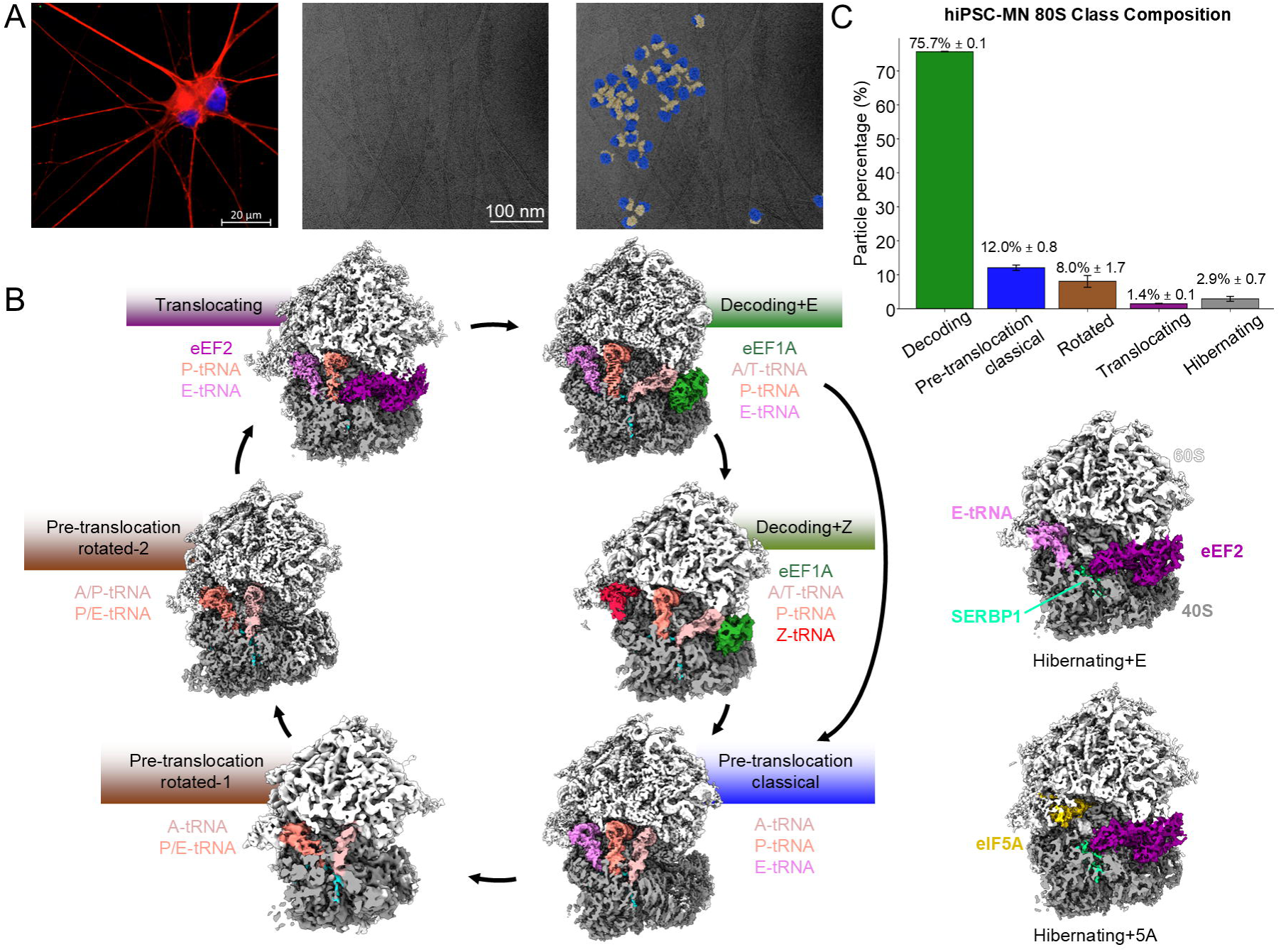
Immunocytochemistry and *in extracto* cryo-EM of hiPSC-MN. (A - left) Combined channels of a representative image of TUJ1 (class III β-tubulin) (red), CC3 (Cleaved Caspase-3) (green), and DAPI (blue). See *SI Appendix*, Fig. 3 for individual channels of antibodies used in this paper. (A - middle) Image of hiPSC- MN lysate with the corresponding overlay (A - right). (B) Cryo-EM maps of 80S states arranged as an elongation cycle (left) and two subclasses of hibernating ribosomes (right) detected in hiPSC-MN lysates. (C) 80S class abundances in cell lines M2 and M4 (shown as % particles ± SEM).

3D classification revealed that 80S ribosomes in elongation states comprise over 95% of particles in hiPSC-MN lysates (*SI Appendix*, Tables S2 and S4), echoing the predominance of elongation states in intact neurons. Furthermore, high-resolution maps help visualize interactions within elongating ribosomes that facilitate the recruitment of the translational GTPases, mRNA decoding and uL16 interactions next to the peptidyl transferase center, as described below. The maps account for the key states of the translation elongation cycle, as in recent analyses of translation in immortalized cell cultures (17, 38). Notably, analyses of motor neurons derived from another hiPSC line of a different individual resulted in nearly identical ribosome state distributions, underlining the reproducibility of *in extracto* cryo-EM classifications (*SI Appendix*, Tables S1 and S2).

The elongation cycle begins when the ternary complex of eEF1A with GTP and an aminoacylated tRNA is delivered to the 80S ribosome. In our dataset, these “decoding” or “codon-sampling” ribosomes are the most abundant state, resembling the cellular populations in other cell types (20, 42). The predominant decoding ribosome class contains tRNAs in the A/T state, P (peptidyl), and E (exit) sites (68.4% of all particles), whereas an additional state contains the “exiting” tRNA in the Z conformation (43), accounting for 7.5% of particles (Fig. 2*B* and *SI Appendix*, Table S4). The high resolution of the decoding center (DC) on the small subunit brings insights into the initial stages of decoding, as discussed in a section below.

In the next step of the elongation cycle, a cognate (complementary to the A-site mRNA codon) aminoacyl-tRNA is accommodated into the A site and is available for peptidyl transfer from peptidyl-tRNA in the P site. In these post-accommodation states, the exit tRNA is positioned in the E or Z site, resembling those in other cell types visualized by *in situ* cryo-EM/ET (38), together accounting for ∼10% of all 80S particles (Fig. 2*B* and *SI Appendix*, Table S4). A fraction of these ribosomes contains lower-resolution density that resembles extended eEF1A•GDP next to the accommodated A-site tRNA (*SI Appendix*, Fig. S4 *L*), likely representing the late steps of aminoacyl-tRNA accommodation (14, 17).

Upon peptidyl transfer from the P- to A-site tRNA, the small ribosomal subunit spontaneously rotates relative to the large subunit, coincident with the movement of the deacylated tRNA acceptor arm into the E site on the large subunit (10, 11, 44, 45), while its anticodon stem loop (ASL) remains in the P site of the small subunit (hence, P/E tRNA). A minor fraction of elongating ribosomes in our dataset features tRNAs occupying the P/E and A/A (i.e., peptidyl-tRNA in the A sites of both subunits), accounting for ∼1% of ribosome particles. The more abundant rotated pre-translocation ribosome contains both tRNAs in hybrid states, P/E and A/P (9.2%, Figs. 2*B* and *C*, and *SI Appendix*, Table S4), a distribution resembling that in other cell types (17). In the subsequent step, elongation factor eEF2 catalyzes translocation of the ASLs on the small subunit, resulting in a ∼1% fraction of near-post-translocation non-rotated ribosomes with eEF2 and tRNAs in the P and E sites (Fig. 2*B*).

Hibernating 80S ribosomes (i.e., lacking mRNA density) comprise less than 5% of particles (Fig. 2*C* and *SI Appendix*, Table S4). Echoing our and others’ observations (14, 38), the predominant hibernating ribosome populations feature eEF2 in its canonical position near the A site, whereas the mRNA tunnel is occupied by SERBP1. The two major classes differ by the occupancy of the E site: one class contains deacyl-tRNA (1.5%), the second features eIF5A (1%; Fig. 2*B* and *SI Appendix*, Table S4). Proteins eEF2, eIF5A, and SERBP1 bind the key functional centers of the ribosomes: eEF2 blocks the sarcin-ricin loop (SRL), SERBP1 blocks the P site, and eIF5A blocks the peptidyl transferase center (PTC). The positioning of these proteins therefore sterically occludes factors that may perturb rRNA functional centers, such as nucleases (14). Both hibernating states are consistent with earlier findings that inactive ribosomes in neurons are bound to eEF2 and SERBP1 (34).

To address potential differences in the translational states among neuronal subtypes and organisms, we next performed *in extracto* cryo-EM of mouse primary cortical neurons (PCN). Like the iPSC-derived human motor neurons described above, mouse PCNs contain nearly 90% translationally active ribosomes (Fig. 3 and *SI Appendix*, Table S2). The ∼90/10 distribution between the elongation and hibernation states is remarkably similar to that in rat hippocampal neurons in recent lower-resolution cryo-ET analyses, which resolved eEF1A- and tRNA-bound elongating ribosomes (19). In our samples,∼85% of ribosomes adopt elongation states similar to those in hiPSC-MN described above, and ∼3% of ribosomes feature release factor eRF1 and ABCE1 indicative of ribosomes during translation termination on stop codons (22). The distribution among elongation states resembled that in hiPSC-MN, with decoding ribosomes being most abundant (63.7%), pre-translocation states accounting for 16.7% of particles, and the nearly translocated eEF2-bound complexes representing a rarer elongation state (3.0%) (*SI Appendix*, Table S2). In addition, an unusual class was detected, with density consistent with four tRNAs bound to elongating ribosomes. Specifically, a non-rotated ribosome with three classical-state tRNAs in the A, P, and E sites features density next to the A-tRNA that closely resembles the ternary complex of eEF1A•GTP with aminoacyl-tRNA (Fig. 3 *A-B*). The ternary complex is positioned further away from the A site than in the canonical decoding complex, because the A site is occupied by a tRNA. The position of eEF1A closely resembles that in the 80S ribosome bound with angiogenin in the A site, which interacts with the ternary complex delivered to the nuclease for cleavage (*SI Appendix*, Fig. S5 *D-E*) (46). These observations suggest that eEF1A has an additional binding site on the 40S subunit, which may be sampled on non-rotated ribosomes. The presence of this state may indicate rare non-productive sampling of the ribosome by the ternary complex, in keeping with the high abundance of eEF1A and tRNA in the cell (Fig. 3*C*). In summary, our analyses of ribosomes in hiPSC-derived motor neurons and mouse cortical neurons reveal that neuronal ribosomes adopt predominantly elongation-competent states, suggesting high translation activity.

**Figure 3.**
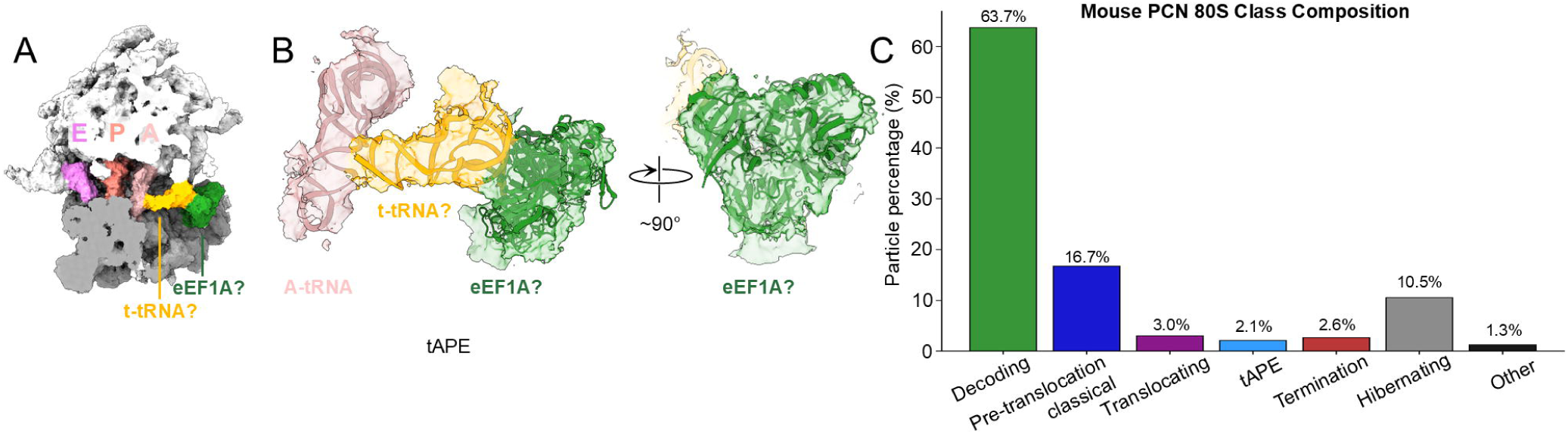
*In extracto* cryo-EM of mouse primary cortical neurons. (A) The overall map and (B) close-up view of the ribosomes with putative A/T, A, P, E-bound tRNAs (tAPE) and eEF1A. (C) 80S class abundances (% particles).

### Low abundance of stalling or collision in neuronal ribosomes and ribosomal subunits

Ribosomes conformations may inform of translational stalling (47, 48), which in neurons is associated with neurodegenerative diseases (6, 8). Ribosomal stalling within cells results in distinct structural features (20). First, ribosome stalling can lead to an up to 20% accumulation of an “off P-tRNA” state, in which the P-site tRNA disengages from the mRNA codon, likely representing a stalled-ribosome disassembly stage. By contrast, such a state, if present, is beyond the detection limit in our neuronal ribosome samples. Similarly, a large General Control of Amino-acid Synthesis 1-like 1 protein (GCN1) accumulates on collided disomes, spanning from the P stalk of the stalled ribosome to the P stalk of the collided ribosome (20, 47). Extensive classifications of the ribosome, including those focused on the P-stalk region described below, did not reveal densities consistent with GCN1 (*SI Appendix*, Fig. S6 and S7). Next, decoding ribosomes with the Z-bound tRNA increase from ∼6% under normal conditions to ∼25% in collision-induced samples, likely due to the limited mobility of the tRNA-supporting L1 stalk, which docks into the neighboring lagging ribosome (20).

Accordingly, stronger density appears next to the L1 stalk, representing a lagging ribosome indicative of ribosome collisions (20). By contrast, the 7.5% population of the Z-bound decoding complex in our dataset and the smaller population of neighboring ribosome density indicate normal translational activity (*SI Appendix*, Fig. S6). Finally, the collided disomes in stressed cells predominantly represent a characteristic configuration of a pioneering non-rotated ribosome with or without a GTPase (such as a DRG protein) followed by a predominantly hybrid-state eEF2-bound rotated ribosome (20). By contrast, ribosomes in intact neurons do not abundantly feature eEF2 (Fig. 1) and the polysome-like configurations represent stretches of eEF1A-bound or hybrid-state ribosomes (Figs. 1*D* and *SI Appendix* Fig. S2 *H*). Collectively, these results are consistent with the low abundance of stalled or collided 80S ribosomes in neurons studied in this work.

The conformations and populations of ribosomal subunits also change upon stress. For example, small 40S ribosomal subunits can undergo restructuring due to eIF2α phosphorylation, which markedly reduces overall initiation (49). Structural analyses of 40S subunit have been more challenging due to the smaller 40S size, resulting in lower resolution that restricts the identification of novel complexes. *In extracto* cryo-EM, however, allows the identification of initiation complexes at near-atomic resolution (26), resolving large populations of 48S complexes that scan the 5’-untranslated regions or bind to the start codons of mRNAs. Furthermore, analyses of immortalized cell lysates revealed eukaryotic initiation factor 1 scanning (eIF1S), also termed eIF5-mimicking protein (5MP), interacting with eIF2 as part of the scanning 48S complex and modulating initiation fidelity (26). Since eIF1S is implicated in neuronal homeostasis and neurodegeneration (50, 51), identification of this protein and other initiation complexes may inform on initiation activity and fidelity in neurons.

To identify the 40S functional states, we used 2DTM with a template comprising the body of the 40S subunit and performed particle classification (*SI Appendix*, Fig. S4 *H-K* and S5). More than half of 48S initiation particles represent three distinct stages of initiation, each comprising ∼20% (Fig. 4*A* and *SI Appendix*, Table S3). During the scanning, the 48S adopts an open conformation, in which the head is tilted away from the body to widen the mRNA tunnel and keep the initiator tRNA in the P site disengaged from mRNA (P_out_) (52–55). In contrast to the *in vitro* assembled complexes, the head-open 48S state in our dataset, in addition to mRNA, P_out_ initiator tRNA and canonical initiation factors eIF1, eIF1A, eIF2, eIF3, contains prominent density in the A site next to eIF1A and the eIF2•tRNA complex. This complex corresponds to the 48S•eIF1S scanning complex (Fig. 4 *A-B*), which we recently identified in an immortalized cell line (26). eIF1S enhances the fidelity of start codon selection (26, 50), and the predominance of eIF1S in the scanning 48S complexes is consistent with the key role of eIF1S (*krasavietz* in Drosophila) in neurodevelopment and memory (56, 57), suggesting extensive regulation of translation initiation on neuronal mRNAs (58). The second 48S state corresponds to the “closed” codon-recognition complex (Fig. 4*A*). Here, upon start codon arrival at the P site, the 48S head tilts toward the body as the initiator tRNA is brought in to base pair with the start codon (P_in_). Finally, a late intermediate that occurs prior to 60S subunit joining features eIF5B bound to the initiator tRNA as eIF2 has been released (Fig. 4*A*) (52). Similar classes have emerged in mouse PCNs (for values see *SI Appendix*, Table S3 and Fig. 4 *C-D*), with strikingly similar particle distributions (e.g., 23% 48S•eIF1S). Together, these findings indicate that 40S subunits extensively interact with the fidelity factor eIF1S and are actively engaged in translation initiation, consistent with efficient translation in neurons.

**Figure 4.**
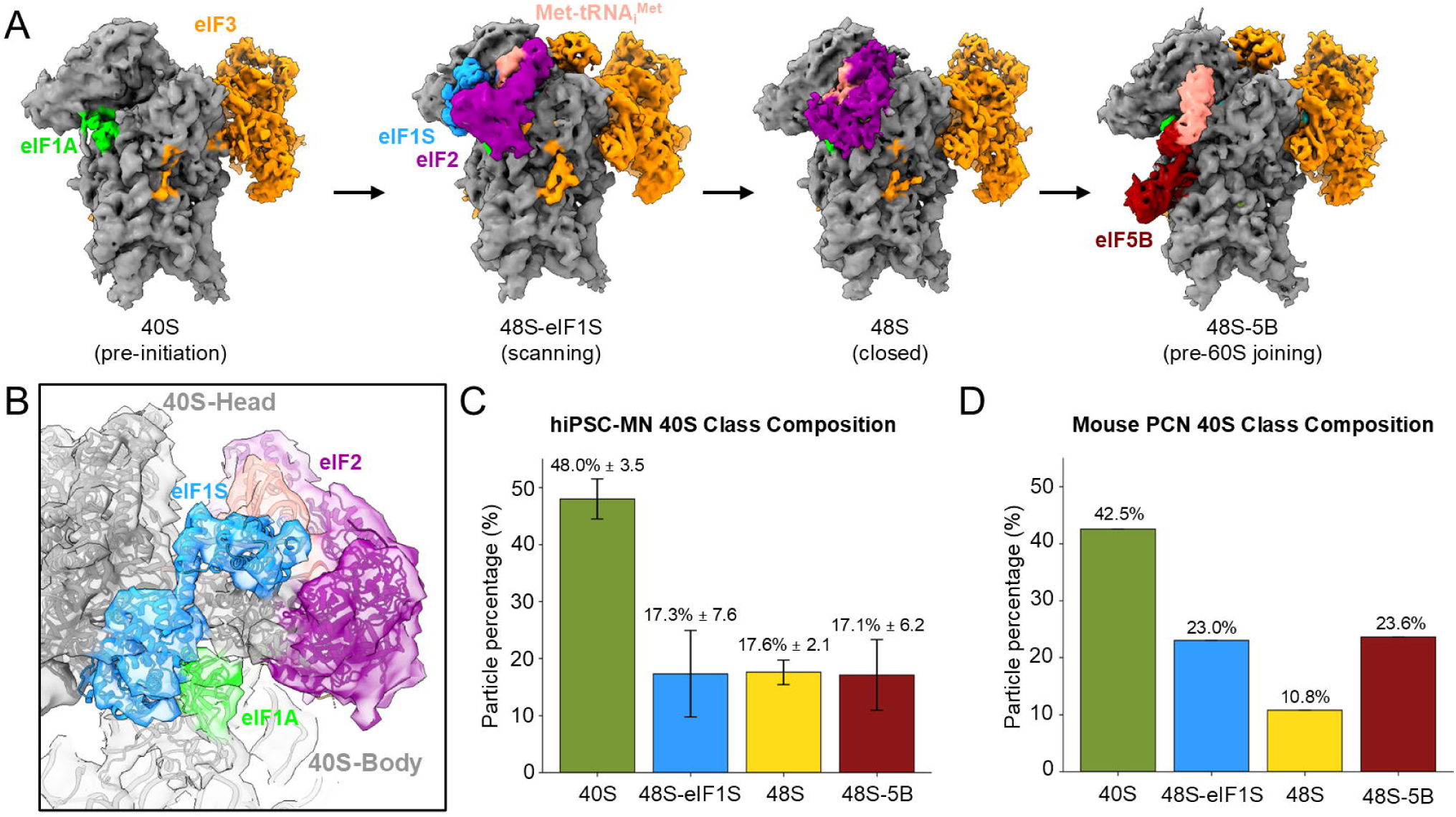
*In extracto* cryo-EM of ribosomal subunits in human iPSC-derived motor neurons and mouse primary cortical neurons. (A) Cryo-EM maps of 40S complexes in hiPSC-MN capturing stages of translation initiation. (B) Close-up view of the 48S-eIF1S map. (C) Class distributions (% particles) in two human iPSC-motor neuron cell lines. (D) Class distributions in mouse primary cortical neurons.

Extensive classification of 60S particles in hiPSC-MN and mouse PCN has not yielded a 60S-tRNA-eIF6-nascent-chain complex indicative of stalling (20). 60S particles from hiPSC-MNs are bound to the anti-association factor eIF6 (*SI Appendix*, Fig. S5 *B*) and represent a small (6.6%) fraction relative to translating 80S ribosomes, indicating that the majority of cytoplasmic subunits are involved in translation. The population of 60S subunits in mouse PCN is also low (2.5%), but in addition to eIF6 they contain the E3 ubiquitin ligase Listerin and tRNA (*SI Appendix*, Fig. S5 *C*), indicating a ribosome-associated quality control (RQC) complex with stalled nascent proteins targeted for degradation (59). Thus, in contrast to hiPSC-MN ribosomes, a minor fraction of ribosomes in mouse PCN appears to experience stalling that is being resolved by the RQC machinery.

### Interactions in the elongating ribosomes: P stalk, DC and PTC

The high resolution of reconstructions of elongating ribosomes provides insights into the roles of interactions at the P-stalk periphery of the ribosome, in the decoding center and in the peptidyl transferase center (Figs. 5 and 6). As eEF1A and other translational GTPases interact with the ribosome, the dynamic P stalk of the ribosome was proposed to interact with these proteins in the vicinity of the ribosomal GTPase center (60, 61) (Fig. 5*A*).

**Figure 5.**
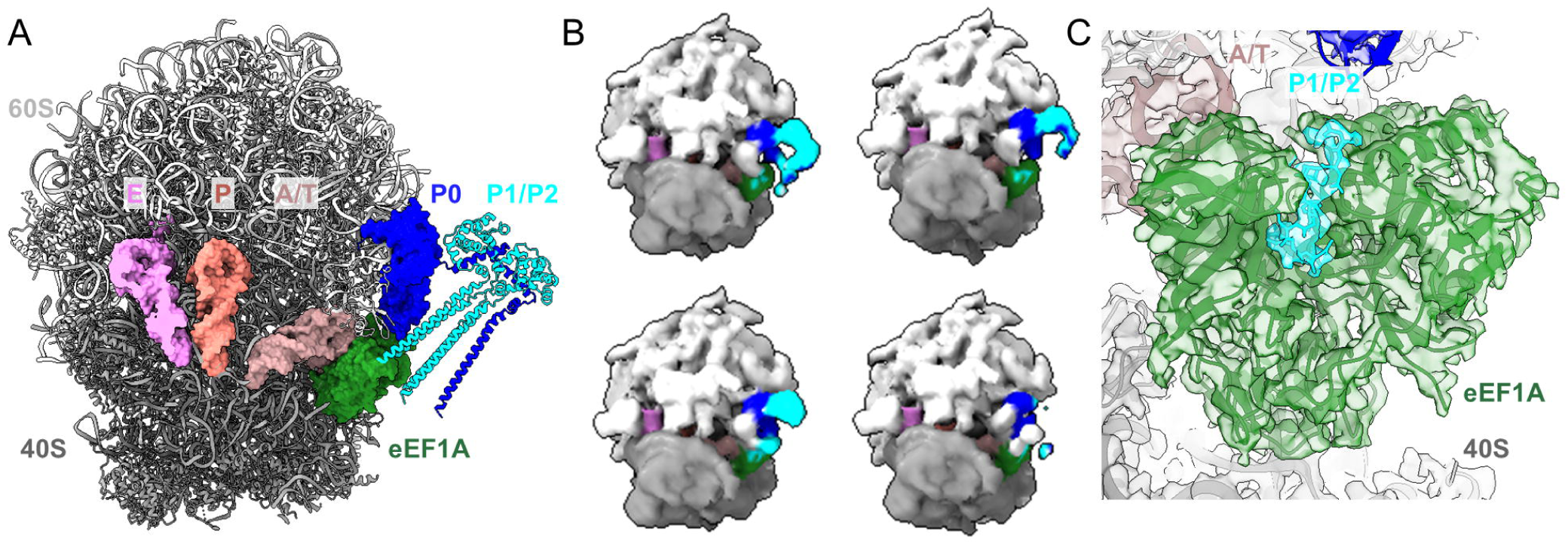
Occupancy and interactions of the ribosomal P stalk. (A) 80S structure of the decoding state (this work) with the P-stalk structure modeled with AlphaFold3 (64). (B) Representative examples of cryo-EM maps demonstrating the dynamics and occupancy of P0 (blue) and P1/P2 (cyan) proteins (see *SI Appendix*, Fig. 5 showing 30 classes resulting from focused classification). (C) The density in the eEF1A pocket is consistent with a P1/P2 interaction.

**Figure 6.**
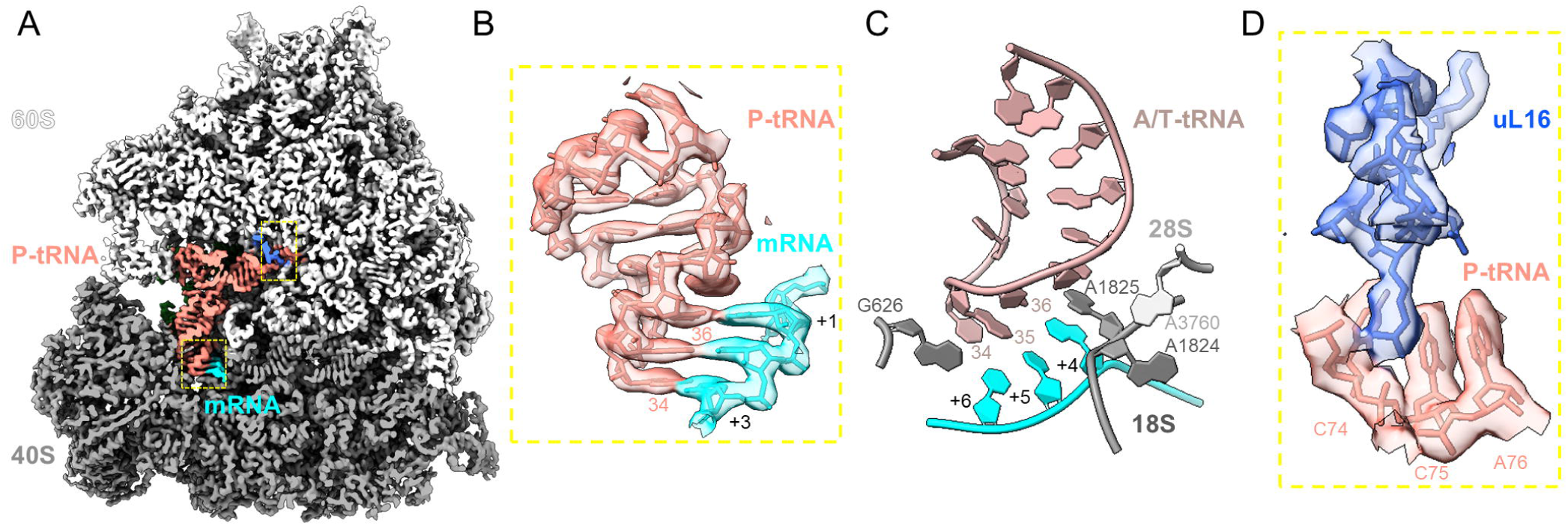
High-resolution features of the decoding 80S ribosomes. (A) Cross-section of the cryo-EM map with P-site tRNA in salmon, mRNA in cyan, 40S in gray, 60S in light gray, and uL16 in blue. The yellow dashed boxes highlight the P-site codon-anticodon interaction and uL16 interaction with the P-tRNA. (B) Close-up view of the P-site codon-anticodon interaction. (C) Interactions in the decoding center (A site) of the ribosome. Also see *SI Appendix*, Fig. 9. (D) Close-up view of the map and model for uL16 interactions with the CCA-end of P-site tRNA in the peptidyl transferase center.

The P stalk is a protein heteropentamer composed of one P0 (also termed RPLP0 and uL10) and two copies each of P1 and P2 (RPLP1 and RPLP2), which share high sequence similarity (62). P0, attached to the 60S subunit, binds P1 and P2, whose α-helical C-termini are exposed to the solvent (Fig. 5*B* and *SI Appendix*, Figs. S7 and S8). The 2.6 Å map of the decoding 80S•eEF1•aatRNA ribosome from hiPSC-MN features an additional density between domains I and III of eEF1A that cannot be accounted for by eEF1A (Fig. 5*C*, cyan). Because the density closely resembles the interaction between homologous archaeal aEF1A and aP1 complex (63) (*SI Appendix*, Fig. S8), we docked AlphaFold-predicted models of P-stalk-eEF1A interactions (64) into the map. The map closely agrees with the C-termini of P1 or P2 placed near domains I and III of eEF1A (Fig. 5*C*). Here, the hydrophobic pocket of eEF1A, composed of Ile181, Leu138, Leu139, Val435, Val437, and Ile345, accommodates the hydrophobic residues of P1 or P2 (Met108, Phe110, Leu112, and Phe113) (Fig. 5*C* and *SI Appendix*, Fig. S8 *D-E*). Just outside the hydrophobic pocket, an acidic stretch of P1 featuring consecutive aspartate and glutamate residues is positioned next to two arginine residues and two lysine residues of eEF1A (*SI Appendix*, Fig. S8 *E*). The electrostatics-rich interaction echoes that in the bacterial counterpart involving the L7/L12 stalk of the 70S ribosome and elongation factor EF-Tu (31). Whereas this cryo-EM density does not allow us to distinguish between very similar P1 and P2 sequences, it is consistent with the proposed function of the P-stalk of increasing the local concentration of eEF1A to facilitate mRNA decoding (61).

P0 and other P-stalk proteins were proposed to be dynamically exchanged within neuronal ribosomes, potentially leading to ribosomes with substoichiometric P stalks differentially engaging in translation or other processes, *i.e.,* “specialized” ribosomes (65). To investigate the P-stalk substoichiometry in neuronal ribosomes, we performed data classification with a focus mask covering the P-stalk to separate particles based on compositional and/or conformational heterogeneity, as in previously reported studies separating numerous ribosome-binding factors (31, 66–70). Extensive classification into 30 classes revealed distinct positions of the P stalk, consistent with its high mobility (*SI Appendix*, Fig. S7). P0 is clearly visible in all classes (*SI Appendix*, Fig. S7). This analysis indicates that ribosomes with substoichiometric P stalk are very rare, if present, in neuronal ribosomes, which predominantly populate the cell body. This analysis does not rule out P-stalk dissociation or fast exchange in localized areas, such as neuronal processes, as the small population of such ribosomes in the lysed neurons may fall below the detection limit of cryo-EM particle analyses. Further, compositional heterogeneity is possible within hibernating ribosomes, whose population in our samples is low. Nevertheless, P0 binding to maturing ribosomes is thought to be irreversible (71), consistent with the high affinity demonstrated by its retention even upon high-salt washes (72). Therefore, its dissociation, if it occurs in cells, would likely require a dedicated mechanism. In summary, our analyses do not reveal translationally active ribosomes lacking the P stalk, consistent with the essential role of the P stalk for cell viability (73) due to its critical role in ribosome biogenesis (74) and GTPase-binding function during translation (60, 61).

As eEF1A brings an aa-tRNA to the A site of the small ribosomal subunit, the tRNA’s ASL samples the decoding center (DC). It is of critical importance for the ribosome to maintain high decoding fidelity to ensure the accurate expression of genetic information. Structural studies of bacterial ribosomes found that the codon-anticodon complementarity results in the closure of the small-subunit shoulder domain (75), bringing the elongation factor toward the GTPase-activating center on the large subunit (31, 66, 76). GTP hydrolysis results in the GTPase dissociation and the acceptance of the aminoacyl-tRNA, whereas near/non-cognate tRNAs fail to induce the efficient domain closure and GTP hydrolysis. The subunit closure coincides with the movement of DC nucleotides G530, A1492 and A1493 (75), as the latter adenosines dock in the minor groove of the complementary codon-anticodon helix. By contrast, non-complementary codon-anticodon interactions fail to efficiently induce DC rearrangements, thus preventing efficient GTP hydrolysis and non-cognate tRNA accommodation (31, 66). In mammalian ribosomes, the corresponding DC nucleotides G626, A1824 and A1825 similarly dock at the cognate codon-anticodon helix resulting in GTP hydrolysis on eEF1A (77), but the transient interactions resulting in tRNA acceptance or rejection prior to GTPase activation remained unresolved (*SI Appendix*, Fig. S9 B). Our structure captures a pre-activation decoding center, in which the codon-anticodon helix is formed, likely between partially complementary mRNA and tRNAs, while G626 remains disengaged from the codon-anticodon helix (Fig. 6*C* and *SI Appendix*, Fig. S9 *A*). A1825 is stretched between the P and A sites to stack on the A-site mRNA codon, so that its rearrangements into the minor-groove-docked position would require flipping around mRNA nucleotides (*SI Appendix*, Fig. S9 A and C). The elongation factor eEF1A is bound with GTP and placed on the 40S subunit shoulder ∼5 Å away from its GTPase-activated position near the SRL (77, 78), in keeping with an early decoding state. The transition of the ribosome from this state into the GTPase-activated state involves a ∼4 Å movement of the shoulder domain coincident with the closure of the decoding center (*SI Appendix*, Fig. S9 *C*). The early decoding ribosome structure in this work is therefore consistent with the universally conserved mechanism of codon recognition, in which the ribosomal DC and 40S closure play the central roles in bridging tRNA base pairing with GTPase activation.

The second key function of the ribosome is to catalyze peptide-bond formation. The reaction is catalyzed by juxtaposing the peptidyl-tRNA and aminoacyl-tRNA substrates optimally in the P and A sites, respectively, (79) and by proton shuttling involving the 2′-OH of the terminal adenosine of P-tRNA (often numbered A76) (80, 81). The peptidyl transferase center is made of universally conserved ribosomal RNA (82) performing the essential function of substrate positioning. Yet, bacterial ribosomal protein bL27 protrudes toward the CCA end of P-tRNA (83–85) (*SI Appendix*, Fig. S10 *A*) and modulates peptidyl transferase activity (86). Our high-resolution cryo-EM maps of the decoding state (Fig. 6 *A-B*) reveal that in eukaryotes, which lack a bL27 homolog, a eukaryote-specific internal loop of uL16 binds the ^74^CCA^76^ end of P-tRNA (Fig. 6*D* and *SI Appendix*, Fig. S10 *B-D*). Here, the guanidinium side chain of Arg110 is coordinated at the phosphates of C74 and C75, as the side chain stacks on ribosomal A4548 (*SI Appendix*, Fig. S10 *B-C*). The backbone carbonyl of Arg110 is positioned to hydrogen-bond with the N4 amino group of C74 (Fig. 6*D* and *SI Appendix*, Fig. S10 *B*). This “P-site loop” of uL16 is positioned similarly in the ribosome with the accommodated A-site tRNA (*SI Appendix*, Fig. S10 *C*), echoing lower-resolution observations in yeast ribosomes (87). Our maps therefore demonstrate how the “P-site loop” of uL16 can contribute to P-site tRNA substrate stabilization in the PTC prior to and after the arrival of the A-site substrate. These findings provide the structural basis for the important role of uL16 loop in translation, as demonstrated by peptidyl transfer inhibition by mutations disrupting this loop (88). Notably, the loop adopts a different conformation in an initiation 80S complex, where it stabilizes the CCA end of the tRNA about to be docked into the peptidyl transferase center upon dissociation of initiation factor eIF5B (PDB: 6WOO) (89), consistent with the important role of this dynamic loop in P-site tRNA stabilization (*SI Appendix*, Fig. S11).

In pre-translocation rotated ribosomes, the uL16 loop is not fully resolved, and its position is occupied by the hybrid-state A/P tRNA. This suggests that the loop becomes destabilized to prevent a steric clash with the translocating tRNA. By contrast, the globular region of uL16 binds near the T stem of A/P tRNA, likely stabilizing this transient state, while P-tRNA is positioned further away (*SI Appendix*, Fig. S12). The rearrangements of the P-site loop of uL16 in our structure are consistent with mutagenesis analyses demonstrating the loop contributes to the intersubunit rotation of ribosomal subunits critical for tRNA translocation (88).

## Conclusions

In this work, we visualized the functional states of ribosomes in neurons using both *in situ* and *in extracto* cryo-EM. In motor neurons differentiated from human iPSCs and in mouse primary cortical neurons, the vast majority of ribosomes are in elongation states, whereas only small fractions account for stalled or non-translating hibernating ribosomes. The ≥90% abundance of elongating cytoplasmic ribosomes in neurons exceeds their abundance in immortalized cell lines (65-80%) that boast very high translation activity (14, 20, 38, 42). This finding emphasizes the exceptional dependence of neurons on translation, consistent with its critical role in neurodevelopment, memory formation and other aspects of brain function. Consistent with mRNA decoding being the rate limiting step of elongation, the majority of ribosomes sample the eEF1A•GTP•aa-tRNA ternary complex to select for the tRNA complementary to the A site codon (Fig. 2). Our 2.6 Å resolution reconstruction of this state revealed the interaction of eEF1A with the mobile P stalk, similar to its archaeal counterparts, stressing the evolutionary conservation of the P-stalk interaction with translational GTPases (63). Consistent with the high affinity of P0 to the ribosome and the essential role of the P stalk in ribosome biogenesis and function, nearly all neuronal ribosomes are equipped with P0, the core P-stalk protein (Fig. 5). The proposed P-stalk exchange among ribosomes (65) is below the level of detection and, if it occurs, may be limited to specific cell locations and likely requires a specific mechanism to be identified. In addition, our structure captures an early decoding step, in which the DC nucleotides interact with the codon-anticodon helix while eEF1A is kept away from the GTPase activation center, preventing GTP hydrolysis during transient sampling of tRNAs in the A site (Fig. 6). The similarity with the structural rearrangements in bacterial ribosomes underscores the universal conservation of the decoding mechanism. Finally, our high-resolution analyses identify a protein-RNA interaction at the peptidyl transferase center. Here, the loop of uL16, present in archaea and eukaryotes, binds the CCA end of the P-site tRNA, in keeping with the role of this loop in peptidyl transferase activity and translocation (88).

The structural composition of ribosomes in neurons is similar to that in other cell types, including the hibernating ribosomes predominantly featuring elongation factor eEF2 (14, 38) and scanning initiation 48S complexes featuring eIF1S (5MP) (26). The nearly complete occupancy of the stringency factor eIF1S on the scanning (head-open) 48S in iPSC-MN and primary cortical neurons (Fig. 4) suggests tight regulation of translation initiation, which remains to be evaluated in future studies. In this work, the predominance of elongation states is consistent with the high basal translational activity in “young” iPSC-derived neurons and fetus-derived cortical neurons. Yet, neurons can achieve different translation activities depending on the subtype, developmental stage, excitation/inhibition status, nutrient availability, stress and other factors (33, 34, 90–92). Our findings emphasize the applicability of 2DTM and *in extracto* cryo-EM in studying translation, including the identification of novel complexes and high-resolution interactions, encouraging future work to elucidate the effects of different factors on translational activity of neurons and other cells.

## Data availability

The structural model for the decoding 80S-eEF1A ribosome from human iPSC-derived neurons, with modeled uL16 P-site loop and partial SERBP1 and P-stalk proteins, has been deposited in the RCSB Protein Data Bank under code 39OL, and the related cryo-EM map used to generate this model has been deposited in the Electron Microscopy Data Bank under code EMD-79652.

## Acknowledgments

We thank Chen Xu, Kangkang Song and Christna Ouch for support with cryo-EM data collection at UMass Chan Medical School; and members of the Grigorieff and Korostelev laboratories for constructive feedback on this study. N.G. is an investigator of the Howard Hughes Medical Institute. This study was supported by the UMass Chan Training Program TL1TR001454 (awarded to S.D.) and US National Institutes of Health grants R21AG084170 (to F.B.G. and A.A.K.), R01NS101986 (to F.B.G.), R01NS057553 (to F.B.G.), R21MH134127 (to A.A.K), and R35GM127094 (to A.A.K.).

## Author contributions

**S.D.** prepared and analyzed cellular and cryo-EM samples; collected and analyzed cryo-EM data; prepared illustrations and drafted the manuscript. **J.E.** assisted with cryo-EM data analyses. **Y.Z., S.L. and D.H.** prepared cells. **X.Z.** and **Z.S.** assisted with cell lysate preparation. **F.-B.G.** designed research, supervised cell sample preparation, and secured funding. **N.G.** designed research, supervised data analysis, drafted the manuscript and secured funding. **A.A.K.** designed research, supervised data analysis, drafted the manuscript and secured funding. All authors revised the manuscript.

