## Supplementary Information for "High-resolution cryo-EM of ribosomes in mammalian neurons"

*Supplementary Index:*

Tables S1-S5

Materials and Methods

Supplementary References

Supplementary Figure Legends

**Table S1:** hiPSC cell lines used in this work

| Cell line | Source | Name in Manuscript |
|---|---|---|
| Human: Isogenic control line 27M91 | Lopez-Gonzalez et al. (2019) | M4 |
| Human: Isogenic control line 26Z90 | Lopez-Gonzalez et al. (2019) | M2 |

**Table S2:** Particle percentages corresponding to 80S classes (translation states) in lysate datasets

**80S Pooled Processing (%)**

| | hiPSC-MN | | Mouse PCN |
|---|---|---|---|
| | M4 Unpooled | M2 Unpooled | |
| Decoding | 75.6 | 75.8 | 63.7 |
| Pre-translocation classical | 11.3 | 12.8 | 16.7 |
| Rotated | 9.7 | 6.3 | |
| Translocating | 1.3 | 1.6 | 3.0 |
| Hibernating | 2.1 | 3.6 | 10.5 |
| Terminating | | | 2.6 |
| tAPE | | | 2.1 |
| Mixed/other | | | 1.3 |

**Table S3:** Particle percentages corresponding to 40S classes (initiation states) in lysate datasets

**40S Pooled Processing (%)**

| | hiPSC-MN | | Mouse PCN |
|---|---|---|---|
| | M4 Unpooled | M2 Unpooled | |
| 40S Pre-IC | 51.5 | 44.5 | 42.5 |
| 48S-5MP | 9.7 | 24.9 | 23.0 |
| 48S (closed) | 15.4 | 19.7 | 10.8 |
| 48S-5B | 23.3 | 10.9 | 23.6 |

**Table S4:** Particle percentages and 40S rotation† in 80S classes from hiPSC-MN M4

| 80S particles (%) | | 40S-Body rotation (°) | 40S-Head swivel (°) |
|---|---|---|---|
| Decoding+E | 68.4 | -1.9 | 0.7 |
| Decoding+Z | 7.5 | -1.8 | 0.7 |
| Pre-translocation classical non-rotated* | 11.4 | 0 | 0 |
| Pre-translocation rotated-1 | 0.6 | 5.8 | 3.2 |
| Pre-translocation rotated-2 | 9.2 | 9.7 | 1.5 |
| Translocation | 0.5 | 1.3 | 16.6 |
| Hibernating+5A | 1 | 5.4 | 14.8 |
| Hibernating+E | 1.5 | 4.6 | 14.8 |

† The rotations were measured relative to the classical-state non-rotated 80S with three tRNAs.

* The classical non-rotated classes include lower resolution maps with putative extended-form eEF1A (*SI Appendix*, Fig. S4*L*)

**Table S5:** Cryo-EM data collection, refinement, and validation statistics for the 80S•eEF1A•GTP•aa-tRNA complex

| | 80S•eEF1A•GTP•aa-tRNA (hiPSC-MN) |
|---|---|
| EMDB Accession ID | 79652 |
| PDB ID | 39OL |
| **Data collection and processing** | |
| Magnification | 81,000x |
| Voltage (kV) | 300 |
| Electron exposure ($e^-$/Å$^2$) | 30 |
| Defocus range (μm) | 0.5-2.0 |
| Pixel size (Å) | 0.83 |
| Number of particles | 368271 |
| **Refinement** | |
| Initial model used (PDB code) | 5LZS |
| Model resolution (Å) | 2.6 |
| FSC threshold | 0.143 |
| Correlation Coefficient (cc_mask) | 0.90 |
| Map-sharpening B-factor (Å$^2$) | -40 |
| Model composition | |
| Non-hydrogen atoms | 226,245 |
| Protein residues | 12,592 |
| RNA residues | 5,830 |
| Average B factor (Å$^2$) | |
| Protein | 55.6 |
| RNA | 62.0 |
| R.M.S deviations | |
| Bond length (Å) | 0.016 |
| Bond angles (°) | 1.030 |
| Validation | |
| MolProbity score | 2.07 |
| Clash score | 3.17 |

| | |
|---|---|
| Rotamer outlier (%) | 7.83 |
| Ramachandran plot | |
| Favored (%) | 95.86 |
| Allowed (%) | 4.10 |
| Disallowed (%) | 0.04 |
| RNA validation | |
| Average suiteness (MolProbity) | 0.496 |

#### Materials and Methods

*iPSC-Motor neuron (hiPSC-MN) culture*

The use of human iPSC lines was approved by the University of Massachusetts Chan Medical School Institutional Biosafety Committee (Docket # I-435-20). Motor neurons were differentiated as described before (1). Briefly, at 70–80% confluency, iPSCs were washed with DPBS (Corning) and incubated in Accutase (Millipore) for 1 min at room temperature. iPSC colonies were washed and scraped with a cell lifter, then plated and expanded in mTeSR1 media (Stem Cell Technologies) in Matrigel (Corning)-coated wells. Twenty-four hours after plating, the culture media was replaced with neuroepithelial progenitor (NEP) media, DMEM/F12, Neurobasal media at 1:1, 0.5x N-2 (Gibco), 0.5x B-27 (Gibco), 0.1 mM ascorbic acid (Sigma), 1X GlutaMAX (Invitrogen), 3 μM CHIR99021 (Tocris Bioscience), 2 μM DMH1 (Tocris Bioscience), and 2 μM SB431542 (Stemgent). After 6-7 days, NEPs were dissociated with Accutase, split 1:6 into Matrigel-coated wells, and cultured for 6-7 days in motor neuron progenitor induction media (NEP with 0.1 μM retinoic acid and 0.5 μM purmorphamine, both from Stemgent). Motor neuron progenitors were dissociated with Accutase to generate suspension neurosphere cultures. After 6 days, the neurospheres were dissociated into single cells by pure Accutase, plated on poly-L-lysine (PLL)/laminin-precoated plates/Cryo-EM grids (carbon side up) in motor neuron differentiation media containing 0.5 μM retinoic acid, 0.1 μM purmorphamine, and 0.1 μM compound E (Calbiochem), 10 ng/mL GDNF (R&D Systems), 10 ng/mL BDNF (R&D Systems), and 1 μg/mL laminin (Sigma) for 2 weeks, and then in the same media without compound E for 4 weeks.

*hiPSC-MN immunofluorescence*

We examined neuronal ribosomes after maturation. Motor neurons were fixed in 4% paraformaldehyde for 15 min and permeabilized with 0.1% Triton X-100 for 15 min. Cells were blocked with 3% BSA for 60 min and incubated with primary antibodies overnight at 4°C. The following primary antibodies were used: mouse anti-MAP2 (1:500, Sigma-Aldrich, Cat# M9942, RRID:AB_477256), mouse anti-tubulin βIII/TUJ1 (1:1000,

Promega, Cat# G7121, RRID:AB_430874), goat anti-ChAT (1:200, Millipore, Cat# AB144P, RRID:AB_90661), rabbit anti-NeuN (1:200, Millipore, Cat# ABN78, RRID: AB_10807945), mouse anti-HSP70 (1:200, Thermo Fisher Scientific, Cat# MA3-007, RRID: AB_325455), and rabbit anti-cleaved caspase 3 (1:200, Cell Signaling Technologies, Cat# 9661S, RRID: AB_2341188). After incubation with the primary antibodies, neurons were washed three times with DPBS, incubated with Alexa Fluor conjugated secondary antibodies (Invitrogen, 1:1,000) for 1 hour at room temperature, and mounted with ProLong Gold Antifade Mountant with DAPI (Thermo Fisher Scientific). Confocal images were acquired with a Zeiss LSM 800 laser scanning confocal microscope and processed with Zeiss ZEN software. See Supplemental Figure 3 for individual channels.

*Grid preparation for lamellae*

Quantifoil $SiO_2$ Au200, R2/2 grids were treated with ethyl acetate by submerging the grids in a glass dish. Once evaporated, another round of treatment was performed overnight. The grids were then glow-discharged on both sides using 25 mA of current with negative polarity for 45 seconds. Afterwards, grids were subjected to UV treatment for 90 min. Grids were incubated overnight at 37°C and 5% $CO_2$ in a PLL solution inside glass dishes of the kind used in (2).

*Staining, plunge-freezing, and microscopy for hiPSC-MN Lamellae*

Before freezing, grids containing hiPSC-MN were incubated with propidium iodide (PI) and Hoechst 33342 (Hoechst) by two methods, in order to ascertain which neurons were viable. In the first method, PI and Hoechst dyes were added directly to the plates containing hiPSC-MN grids. The resulting media was pipetted and swirled gently before being incubated for 3-5 min, followed by plunge-freezing of the grids (settings below). In the more extended method, the media was removed from the dishes containing hiPSC-MN before adding 2 mL of 37°C DPBS. The media was then gently swirled and removed before adding DPBS again. The DPBS was again removed and a mixture of 5 mL warm media with 5 µL PI and 5 µL Hoechst dyes was added. When stain was used, it was added to the media (37°C) and mixed well before use. After incubating for 3-4 min, suction was used to remove the stain and media. The plates were then washed once with DPBS before adding 2-3 mL of media without stain.

We did not observe noticeable differences between grids prepared using these two methods. Grids were loaded onto a Leica EM GP2 plunger, back-side blotted at 37°C for 8 seconds in a 95% humidity environment, plunge-frozen in liquid ethane at -184°C and stored in liquid nitrogen until focused ion beam (FIB)-milling. Milling target selection was aided by the iFLM Correlative System (Thermo Fisher Scientific) mounted on an Aquilos 2 Cryo-FIB. Channel wavelengths for PI and Hoechst dyes were used according to manufacturer guidelines. Areas with a high signal corresponding to Hoechst and a low (or absent) PI signal were selected as targets for FIB-milling. FIB-milling and DeCo-LACE imaging were carried out as in (3).

*Cryo-EM data processing and 2DTM for hiPSC-MN Lamellae:*

hiPSC-MN lamella were imaged on a 300 kV Titan Krios electron microscope equipped with a K3 direct electron detector. Movies were collected in super-resolution mode at a pixel size of 0.53 Å. Each movie consisted of 30 frames with a total exposure of approximately 30 $e^-$/Å$^2$. Movies were processed using the DeCo-LACE implementation (github.com/jojoelfe/decolace) in *cis*TEM (4). Motion correction was performed using unblur (4) and micrographs were binned to a final pixel size of 2 Å. CTF parameters were estimated using CTFFIND4 (5). To perform 2DTM we generated a 60S template from PDB 6QZP using the simulate program (6) in *cis*TEM, with the center flag on, B-factor scaling set to 1, a box size of 320 x 320 pixels, and a pixel size of 2 Å. Before template generation, chains for the 40S subunit, mRNA and tRNA were manually deleted from the PDB file. Orientations and defocus of detected targets were refined using a GPU-accelerated version of the match_template program in *cis*TEM (7), with a 3 degree out-of-plane and 2 degree in-plane step size, and a defocus range of 900 Å with a step size of 300 Å.

Using *cis*TEM, detected targets were extracted from the micrographs with a box size of 320 x 320 pixels. We calculated 3D reconstructions from the detected targets, followed by refinement and classification. We excluded targets based on distance to the beam edge (25 Å), pixel variance (0.3), and 2DTM SNR (7.1). 3D refinement was completed in the *cis*TEM GUI using the 2DTM template as a reference (8). Briefly, one round of x, y shifts, followed by one round of angular refinement were performed at Nyquist (4 Å). The particle stacks and refinement parameters from all lamellae were exported from *cis*TEM and combined into a single image stack and set of parameters. The particle images were binned further to a pixel size of 4 Å for further processing using Frealign v9.11 (9). One round of x, y refinement was performed against a 3D reconstruction calculated from the exported stack and parameters using low and high-resolution limits of 50 Å and 16 Å, respectively. The low-resolution limit for any other refinement steps was set to the default 300 Å, unless otherwise stated. 3D classification was done using a focus mask applied to an area comprising the GTPase and the A/T tRNA binding sites (coordinates 397.5 Å, 253.2 Å, 351.3 Å, with a radius of 65 Å). 100 rounds of classification were run with a high-resolution limit for classification set to 10 Å. Particles were classified into 24 classes with the parameter_mask set to “0 0 0 0 0” (only the occupancy was refined). Data collection, 2DTM, and montage creation were completed according to ref (2). The overlay was rendered using the Molecular Nodes package in Blender (10.5281/zenodo.6540846, github.com/BradyAJohnston/MolecularNodes).

*Semi-permeabilization and cytoplasm extraction for hiPSC-MN*

Terminally differentiated cell cultures were semi-permeabilized in the same manner regardless of cell line. Briefly, media was removed from each of the wells of the 6-well plate before gently washing with 2-4 mL of warm DPBS. DPBS was then removed and Accutase was added to remove the adherent hiPSC-MN from the coated dish. Cells were gently swirled before being detached and pipetted into a 15-mL tube and spun down at room temperature for 2 min at 100 g. The resulting supernatant was removed before adding 100-400 µL of semi-permeabilization buffer (25mM HEPES-KOH, 110mM KoAC, 15mM Mg(oAC)$_2$, 1mM DTT, 0.015% Digitonin, 1x Protease Inhibitor (Sigma),

0.5x RNase In (Promega), 1mM EGTA). This mixture was left on ice for 4 min and then at room temperature for 1 min, and finally spun down at 800-2500 g for 3-5 min. The supernatant was transferred to a new tube before measuring the absorbance at 260-nm.

*Plunge-freezing of hiPSC-MN extracts*

Quantifoil Au200, R2/2 grids were glow discharged on both sides using a current of 25 mA with negative polarity for 45 seconds. 2-5 µL of sample was applied to the grids with 10-60 seconds of contact time before back-side blotting. Grids that contacted blotting paper but did not successfully blot had 2-5 µL of buffer (25mM HEPES-KOH, 110mM KoAC, 15mM $Mg(oAC)_2$, 1mM DTT, 0.015% Digitonin, 1x Protease Inhibitor (Sigma), 0.5x RNase In (Promega), 1mM EGTA) added to the back of the grid to aid blotting. Blotting lasted for 8 seconds at 4°C and 95% humidity using a Leica EM GP2 plunger. Grids were plunge-frozen in liquid ethane at -184°C and stored in liquid nitrogen.

*Cryo-EM data collection and image processing for hiPSC-MN extracts*

hiPSC-MN extractions were imaged on a 300 kV Titan Krios electron microscope equipped with a K3 direct electron detector with a target underfocus of 0.5-2.0 µm. SerialEM was used to automate data collection (10). Movies were collected in super-resolution mode at a pixel size of 0.53 Å. Each movie consisted of 30 frames with a total exposure of approximately 30 $e^-$/Å$^2$ in each dataset. Movies were motion-corrected using MotionCor2 (11) using the -FtBin, -Patch (7 5), and -Gain options. FtBin = 3 (1.59 Å/pixel) was used for the 2DTM search, and FtBin = 2 (1.06 Å/pixel) was used for particle extraction and refinement. The only exception is that the physical pixel size of 1.06 Å was used for the 40S-Body in cell line M2 extract data processing. Motion-corrected images were imported into *cis*TEM and the image defocus was estimated using CTFFIND4 (5).

*2DTM and classification of hiPSC-MN extract data*

As previously described (6), we generated a 60S template from PDB 6QZP using the simulate program, after removing the chains for the 40S subunit, mRNA and tRNA from the PDB file. We chose a box size of 512 x 512 pixels, and a pixel size of 1.59 Å to match the pixel size of the micrographs. 2DTM was performed with a 3.5 degree out-of-plane and 2.5 degree in-plane step size, while the defocus search was turned off. Detected targets were extracted from 2x binned micrographs (pixel size 1.06 Å) with a box size of 512 x 512 pixels. Particles from all hiPSC-MN datasets were pooled into a single particle stack and manually refined against a 3D reconstruction. To obtain the highest-resolution reconstruction of the decoding 80S•eEF1A•GTP•aa-tRNA complex, additional hiPSC-MN datasets (beyond M2 and M4, including different media and treatments) were pooled, which were not used to obtain particle distributions in M2 and M4 reported here. Independent classification of the M2 and M4 pool resulted in similar particle distributions, underscoring the consistency of the classification approach. Refinement consisted of one round of x, y shifts, one round of angular refinement, and one round of defocus and beam tilt refinement all with a high-resolution limit of 5 Å. For 3D classification in Frealign version 9.11 (9), particle stacks were binned 8x to a pixel size of 8.48 Å.

*80S classification for cell line M4*

The final reconstruction before any classification had an FSC-estimated resolution of 2.7 Å. Using a focus mask comprising the GTPase and A/T tRNA binding sites (coordinates 338.9 Å, 193.9 Å, 314.8 Å, with a radius of 80 Å), 100 rounds of classification were run with a high-resolution limit for classification set to 17 Å in Frealign, as described above, into 50 classes with the parameter_mask set to "0 0 0 0 0" (only the occupancy was refined). Classes consistent with decoding ribosomes were pooled for sub-classification. Class membership was based on an occupancy > 50%.

To identify the occupancies of the E/Z sites, another 100 rounds of classification were run as before, except specifying 30 classes and using a focus mask on the E and Z site tRNA (coordinates 333.3 Å, 354.9 Å, 351.0 Å, with a radius of 85 Å). The same classification parameters and focus mask were used on pooled non-rotated ribosomes. For pooled eEF2 containing ribosomes, 8 classes were specified, and a focus mask covering the P-site tRNA (coordinates 327.4 Å, 291.8 Å, 293.9 Å, with a radius of 35 Å) was used. See Supplemental Figure 4 for breakdown of sub-classification of 80S particles.

*80S classification for cell line M2*

To determine if our class distributions were reproducible, we repeated our *in extracto* experiments with a second cell line (cell line M2 – *SI Appendix,* Table S1). Data collection was completed in the same manner as before, with the same template and 2DTM parameters. Particles from both datasets were pooled and refined together before classification. After classification, datasets were "unpooled" and their percentages and standard error of means calculated. Our 80S classes showed SEMs ranging from 0.1% to 1.7% (Figure 2C). The most populated classes of decoding states had a mean of 75.7% and an SEM of 0.1% when classifying the datasets together. The remaining particles were assigned to the classical state (12.0% with an SEM of 0.8%), the rotated-2 state (8.0% with an SEM of 1.7%), translocation state (1.4% with an SEM of 0.1%), and the hibernating state (2.9% with an SEM of 0.7%). Despite numerous attempts, particles from cell line M2 could not be sub-classified in the same manner as cell line M4.

The final reconstruction before any classification had an FSC-estimated resolution of 2.7 Å. Using a focus mask comprising the GTPase and A/T tRNA binding sites (coordinates 345.9 Å, 186.4 Å, 317.6 Å, with a radius of 80 Å), 120 rounds of classification were run with a high-resolution limit for classification set to 17 Å. Particles were classified into 40 classes with the parameter_mask set to "0 0 0 0 0" (only the occupancy was refined). Classes consistent with decoding ribosomes were pooled for sub-classification. Class membership was based on an occupancy > 50%. Particles with a cumulative occupancy for a given class (for example, rotated-2) of over 50% were considered to be in the corresponding state of this class. For example, if classes 2, 5, 6, and 7 contained densities corresponding to the rotated-2 state, the occupancies of 2, 5,

6 and 7 would be summed for a given particle. This accounting differs from the frealign_calc_stats program, which sums the occupancy for every particle in each class and then rounds to the nearest integer. After classification, particles from datasets were combined in order to reach a higher resolution for both 80S and 40S classes.

*40S classification of cell lines M2 and M4*

To search for 40S targets, we generated a 40S-Body template from PDB 6QZP using the simulate program as before. This time we removed the chains for the 60S subunit, mRNA and tRNA from the PDB file, as well as the head region of 40S. For cell line M2 we chose a box size of 384 x 384 pixels, and a pixel size of 1.06 Å. For cell line M4 we chose a box size of 320 x 320 pixels, and a pixel size of 1.59 Å. The 2DTM search was run using 3 degrees out-of-plane and 2 degrees in-plane angular step size for cell line M2 and a 3.5 degrees out-of-plane and 2.5 degrees in-plane angular step size for cell line M4, both without searching defocus. Particle stacks from each hiPSC-MN dataset were generated before classifying 40S classes from 80S. Detected targets from both cell lines were subjected to an initial classification individually to parse 40S vs 80S classes using a pixel size of 8.48 Å. Cell line M4 utilized an 80 Å focus mask at the position where eEF1A and A/T tRNA would be positioned. Cell line M2 did not utilize a focus mask. Particles with over a 50% occupancy for 40S classes from both cell lines (M2 and M4) were combined and had an SNR threshold of 7.5 before extracting particles from images. The particle stack was refined against the template for one round as before (x,y shifts, Euler angles, beam tilt and defocus). 40S containing classes from both cell lines were pooled from the 1.06 Å images. The final reconstruction for the 40S pooled stack before sub-classification was estimated at 3.0 Å resolution (FSC = 0.143). Classification of this substack using a focus mask at position 280.9 Å, 335.2 Å, 212.2 Å, and a radius of 50 Å yielded 6 classes (Supplemental Figure 4I-1K). The FSC curves for individual classes are shown in Supplemental Figure 4H. Substacks were “unpooled” to determine class distributions.

*Mouse primary cortical neuron preparation*

All procedures involving mice were approved by the University of Massachusetts Chan Medical School Institutional Animal Care and Use Committee (PROTO202200010). Pregnant BALB/c E16-E18 mice were sacrificed by cervical dislocation and soaked in 75% ethanol for several minutes for surface disinfection. Fetal mice were removed and soaked in 75% ethanol before being rinsed and washed twice with PBS. The cortical hemispheres of mouse brains were extracted and the meninges and vascular tissues were removed. The cerebral cortex was transferred to a 15-mL centrifugation tube and washed twice with pre-cooled PBS. Brain tissue was then digested using ~2 mL of 0.25% trypsin EDTA + DMEM basal media (1:1) + DNase and incubating in a water bath at 37°C for 20 min. After digestion the supernatant was discarded. Shortly after, 3 mL of 4°C pre-cooled seeding media (DMEM/F12 basal media + 10% FBS + 1% penicillin–streptomycin) was added. To fully dissociate the tissue, we gently pipetted up and down ~20 times with a wide-mouthed 1 mL pipette tip, then added an additional 3 mL of the pre-cooled seeding media and pipetted ~20 times. The cell suspension was then filtered with a wet 40 μm filter and centrifuged at 80 g for 5 min. The supernatant was discarded before resuspending the cell pellet with pre-warmed seeding media and performing a cell counting assay. The cell density of the cell suspension was adjusted to $5\times10^4$ cells/mL. Cells were seeded on poly-D-Lysine (PDL)-coated plates for 48 hours before the media was changed to the continuous culture media (Neurobasal media + 1x B-27 + 1x GlutaMAX). Continuous culture media was replaced 1/2-2/3 every other day while the neurons matured.

*Mouse cortical neuron semi-permeabilization and extraction*

Day 16 cell cultures were semi-permeabilized following previously established protocols (12). After briefly washing neuronal cultures with cold PBS, 250 μL of semi-permeabilization buffer (25mM HEPES-KOH, 110mM KoAC, 15mM $Mg(oAC)_2$, 1mM DTT, 0.015% Digitonin, 1x Protease Inhibitor (Sigma), 0.5x RNase In (Promega), 1mM EGTA) was added to each well of a 6-well plate. Cultures were incubated at 4°C for 5 min before letting them stand at room temperature for 1 additional minute. The sample was then collected into a separate tube. Semi-permeabilization, incubation, and collection were repeated once more. The sample was then centrifuged for 5 min at 4°C and 1000 g. Amicon filters were used to concentrate the sample to several hundred ng/µL.

*Plunge-freezing of mouse cytoplasm extractions*

Quantifoil Au200, R2/2 grids were glow discharged on both sides using a current of 25 mA with negative polarity for 45 seconds. 3.5 µL of sample was applied to the grid with 2-5 min of contact time before back-side blotting. Grids that contacted blotting paper but did not successfully blot had 3.5 µL of buffer added to the back of the grid to aid blotting. Blotting lasted for 8 seconds at 4°C, 95% humidity using a Leica EM GP2 plunger. Grids were plunge-frozen in liquid ethane at -184°C and then stored in liquid nitrogen.

*Cryo-EM data collection and image processing of mouse cytoplasm extractions*

Mouse lysate extractions were imaged on a 200 kV Talos electron microscope equipped with a K3 direct electron detector, and a target underfocus of 0.5-2.0 µm. SerialEM was used to automate data collection (10). Movies were collected in super-resolution mode at a pixel size of 0.41 Å. Each movie consisted of 30 frames with an exposure of approximately 30 $e^-$/Å$^2$. Movies were motion-corrected using MotionCor2 using the -FtBin = 3, -Patch (7 5), and -Gain options (11). Motion-corrected images with a pixel size of 1.23 Å were imported into *cis*TEM and the image defocus was estimated using CTFFIND4 (5).

*2DTM and classification of mouse extract data*

To perform 2DTM we generated a 60S template from PDB 7CPU using the simulate program (6) in *cis*TEM, with the center flag on, B-factor scaling set to 1, and a box size of 512 x 512 pixels with a pixel size of 1.23 Å. Before template generation, chains for the 40S subunit, mRNA and tRNA were manually deleted from the PDB file. Orientations and defocus of detected targets were refined using a GPU-accelerated version of the match_template program in *cis*TEM, with a 3.5 degree out-of-plane and 2.5 degree in-plane step size while keeping the defocus constant. Detected targets were extracted from the micrographs using a box size of 512 x 512 pixels. 2DTM targets were refined in *cis*TEM as above, with one round of x, y shifts, one round of angular refinement, and one round of defocus and beam tilt refinement at a high-resolution limit of 5 Å. The final reconstruction yielded a resolution of 2.8 Å (FSC = 0.143) before further classification. Particle images were binned 8x to a pixel size of 9.84 Å and classified using Frealign version 9.11 (9) and a focus mask comprising the GTPase and A/T tRNA binding sites (coordinates 271.6 Å, 411.9 Å, 324.2 Å, with a radius of 80 Å). 100 rounds of classification with the parameter_mask set to “0 0 0 0 0” (keeping alignment parameters constant) were run with a high-resolution limit for classification set to 20 Å, yielding 30 classes. Sub-classification of the tAPE class with a focus mask at 268.3 Å, 418.2 Å, 332.4 Å and a radius of 80 Å was run for 50 additional rounds, yielding 2 classes.

*Structure of the human 80S•eEF1A•GTP•aa-tRNA decoding complex from hiPSC-MN extract*

The structure of the rabbit 80S•eEF1A•GDP•aa-tRNA complex (PDB 5LZS) was used as a starting model to fit ribosomal subunits and other components into the 2.6 Å cryo-EM map using UCSF ChimeraX (13). Novel density for the uL16 P-site loop demonstrates minimal bias against the 60S template, in which this loop was modeled differently (*SI Appendix*, Fig. S13). The structures of human 80S complexes (PDB 8G5Z, 8G60 and 8G61) (14), rabbit 80S•GTPBP1 complex (PDB 9YPG) (15), P1 structure from an AlphaFold3-predicted model of eEF1A (residues 1-237, 257-328, 337-443)•P1 (residues 65-115), GTP (PDB 6JI2) (16), SERBP1 (shown in *SI Appendix*, Fig. S14) from AlphaFold3 and PDB 9AZM, and EBP1 (6Z6N) (17) were used as references to model well-resolved regions using Pymol and UCSF ChimeraX (13, 18), followed by refinements using Isolde (19) and then phenix.real_space_refine v.1.4 (20). Metal ion picking (with phenix.douse v.2.0; ions modeled as $Mg^{2+}$), global minimization, simulated annealing and B-factor (adp) refinements in phenix.real_space_refine yielded a wellfitted 2.6 Å structural model (CC=0.90) with good stereochemical parameters (*SI Appendix,* Table S5).

#### Supplementary Figure Legends

***SI Appendix,* Fig. S1:** Cryo-CLEM-FIB and 2DTM montaging of hiPSC-MN. (A) SEM image taken from an Aquilos 2. (B) SEM image taken at an angle showing an example hiPSC-MN targeted for milling. (C) SEM image of an hiPSC-MN post-milling. (D-F) Cell viability screening of hiPSC-MN on grids. Non-viable cells are represented by the green (PI) signal while viable cells are represented by the blue signal (Hoechst 33342). Blue and green together produce a cyan signal, which indicates non-viable cells. Only cells with blue signal were taken for milling. (G) The full lamella DecoLACE montage from numerous individual micrographs. A smaller area of this image is shown on Fig 1*A*.

***SI Appendix,* Fig. S2:** *In situ* ribosome features. (A) 60S template used for 2DTM. (B) Structure of the decoding ribosome (this work) with RACK1 facing the reader and (C) the 3D reconstruction showing strong density for RACK1. (D) Structure of the decoding ribosome (this work) with EBP1 on the right in orange and (E) the 3D reconstruction showing strong density for EBP1. (F and G) Two low-resolution classes from the lamellae dataset, with eEF1A-like density in forest green (top) and P-site tRNA-like density in salmon (bottom). All structures and reconstructions were generated using ChimeraX (13). (H) Examples of micrograph areas with ribosomes arranged into polysome-like structures, colored according to the occupancy of eEF1A (green) or hybrid-state tRNAs (purple).

***SI Appendix,* Fig. S3:** Individual fluorescence channels from immunocytochemistry assays on hiPSC-MN; (I) red, (II) green, (III) blue, and (IV) merged. (A) Cell stress and post-mitotic neuron identity: HSP70 (AI) and NeuN (AII). (B) Neuron identity and apoptosis: TuJ1 (BI) and CC3 (BII). (C and D) Neuronal identity: (CI and DI) MAP2 and (CII and DII) ChAT. Nuclei were counterstained with DAPI (AIII-DIII).

***SI Appendix,* Fig. S4:** 3D classification schematics for 80S and 40S particles in hiPSC-MN extracts. (A) 3D reconstruction from pooled datasets of 2DTM targets detected using a 60S template, calculated after the refinement of x, y shifts, Euler angles, and defocus. Subsequent classification results in panels B, D, E, F, I, J, and K show the reconstructions in gray. Particle numbers and classes correspond to *SI Appendix,* Table S4 (excluding cell line M2 for 80S classes). If particles were pooled from multiple classes, the number of classes is shown in blue outside of the brackets. Particle percentages are given below each class and are obtained from frealign_calc_stats (21). Weaker density of tRNAs and bound factors are indicated schematically as thinner lines, for example tRNAs in eEF2-containing classes in B. (B) Initial classification results from the pooled dataset. (C) The schematic for ligands bound to the ribosome and other features. (D-F) Sub-classifications of various classes shown in panel B. (G) Fourier Shell Correlation (FSC) plots for reconstructions of 80S and (H) 40S classes. (I and J) 3D reconstructions and classifications from pooled datasets of particles detected by 2DTM using the 40S body template. (K) Pooling of M2 and M4 40S containing particles followed by SNR thresholding and refinement against the 40S-Body template using one round of x, y shifts, Euler angle, and defocus refinement. (L) Extended-form eEF1A-like density is shown in green for states containing E-site and Z-site tRNA.

* Indicates that the occupancy of a single class was taken, and if a given particle had >50% occupancy, it was used for either sub-classification or generating a final stack before generating a 3D reconstruction.

** Indicates that for a given particle the occupancy is summed among all classes that share the same class type. For example, if class 2 and 7 correspond to the same ribosome state, the occupancy for each particle in class 2 and 7 is added together. Values over a 50% occupancy threshold are used for stack generation.

***SI Appendix,* Fig. S5:** *In-extracto* cryo-EM of hiPSC-MN ribosomal subunits and the tAPE state from mouse PCN. (A) The 40S-body template used in this work for 2DTM. (B-C) 60S ribosomal subunit states from hiPSC-MN (B) and mouse PCN (C). (D) The overall map (left) and close-up view of the ribosomes with putative eEF1A•tRNA and A-, P-, and E-site bound tRNAs (tAPE) on the right. (E) The position of eEF1A•tRNA off the A site in an 80S•Angiogenin complex is similar to that in the putative tAPE complex (PDB 9BDP; the surface shows the 80S•Angiogenin structure colored similarly to that in panel D).

***SI Appendix,* Fig. S6:** Classification with a mask over the ribosomal E site separates ribosomes with Z-tRNA (red) and the major classes with E-tRNA (green).

***SI Appendix,* Fig. S7:** Results of a classification of the decoding-state 80S•eEF1A particles with a focus mask covering the P stalk. The 60S is light gray, the 40S is gray, P0 is blue, P1/P2 is cyan, eEF1A is forest green, and E-site tRNA is violet.

***SI Appendix,* Fig. S8:** Putative P-stalk interactions with eEF1A. (A) View of AlphaFold3-predicted P0/P1/P2 structure on the 80S codon sampling class, structures aligned via matchmaker in ChimeraX on the P0 surface bound region (13). The C-terminus of P1 was fit into the density using Isolde (19). (B) Comparison of the crystal structure of archaeal aEF1A•aP1 complex (PDB 6JI2) in pink with AlphaFold3-assisted eEF1A (green) and P1 (cyan) (22). (C) eEF1A, GTP, and P1 colored as a function of predicted Local Distance Differences (pLDDT; the colors are shown in the key to the right). (D) Local areas of hydrophobic interactions between eEF1A and putative P1/P2. (E) Local areas of eEF1A at the interface between domain I and III are rich in basic amino acids (Arg381, Arg382, Lys179, and Lys180), which coincide with the predicted interface of an area rich with acidic amino acids of P1 or P2.

***SI Appendix,* Fig. S9:** Open and closed decoding center (DC) conformations in human 80S•eEF1A decoding complexes with different positions of eEF1A relative to the GTPase-activating sarcin-ricin loop (SRL). (A) Open DC in hiPSC-MN decoding-state ribosomes (this work). (B) Open DC in HEK293T decoding ribosomes (PDB 8G60). Poorly resolved A/T-tRNA residues are colored a lighter shade of salmon. (C) Closed DC in 80S with eEF1A at the SRL (GTPase-activated state; PDB 8G5Z). (D) Cryo-EM density (mesh) of hiPSC-MN ribosomal DC (this work).

***SI Appendix,* Fig. S10:** uL16 interactions across different species and functional states. (A) Bacterial uL16 and bL27 interactions with P-site tRNA in the 70S ribosome. P-site tRNA is orange, bL27 teal, and uL16 purple (PDB 4V5G). (B) hiPSC-MN 80S•eEF1A decoding complex P-site tRNA interactions with the P-site loop of uL16 (this work). (C) Cryo-EM density for the uL16 P-site loop region in the 80S•eEF1A decoding complex with A/T tRNA (left) and 80S with the accommodated A-site tRNA (right). (D) Amino acid sequence comparison of archaea, yeast, human, and mouse uL16. Multiple sequence alignment was performed using Clustal omega 1.2.4 (23); Uniprot codes for individual species are (24): human [Q96L21], mouse [P86048], yeast [P41805], and archaea [Q9YEY5].

***SI Appendix,*** **Fig. S11:** Comparison of uL16 P-site loop and P-site tRNA positions in the yeast initiation 80S•eIF5B complex (PDB 6WOO) and hiPSC-MN 80S•eEF1A decoding complex (this work). (A) Different P-tRNA positions in the initiation and decoding complexes relative to the 60S subunit (alignments were performed via 28S rRNAs). (B) The P-site loop of uL16 interactions with the P-site tRNA in PDB 6WOO and (C) 80S•eEF1A complex (this work). A close-up view in B and C at the bottom of the figure shows distinct positions of Arg110 relative to the tRNA 3′-CCA ends.

***SI Appendix,*** **Fig. S12:** Interactions of the globular domain on uL16 with A/P-tRNA (PDB 6Y57 shown for the rotated-2 state in this work) and P-tRNA (80S•eEF1A decoding state). (A-B) Different positions of the T stem regions of A/P and P-site tRNA relative to uL16. (C-D) Close-up views of uL16 positions relative to A/P-tRNA and P-tRNA.

***SI Appendix,*** **Fig. S13:** The difference in the conformation of the P-site loop of uL16 in the resulting 80S•eEF1•GTP•aa-tRNA map (blue) from that in the 60S template used for 2DTM (gray).

***SI Appendix,*** **Fig. S14:** SERBP1 C-terminal tail interactions with the 80S•eEF1•GTP•aa-tRNA decoding ribosome. (A) Cryo-EM density for the resolved regions of SERBP1 (olive) on the small subunit. (B) The interactions between RACK1 (purple) and the $^{404}$FPxL$^{407}$ C-terminal motif of SERBP1 (olive map).

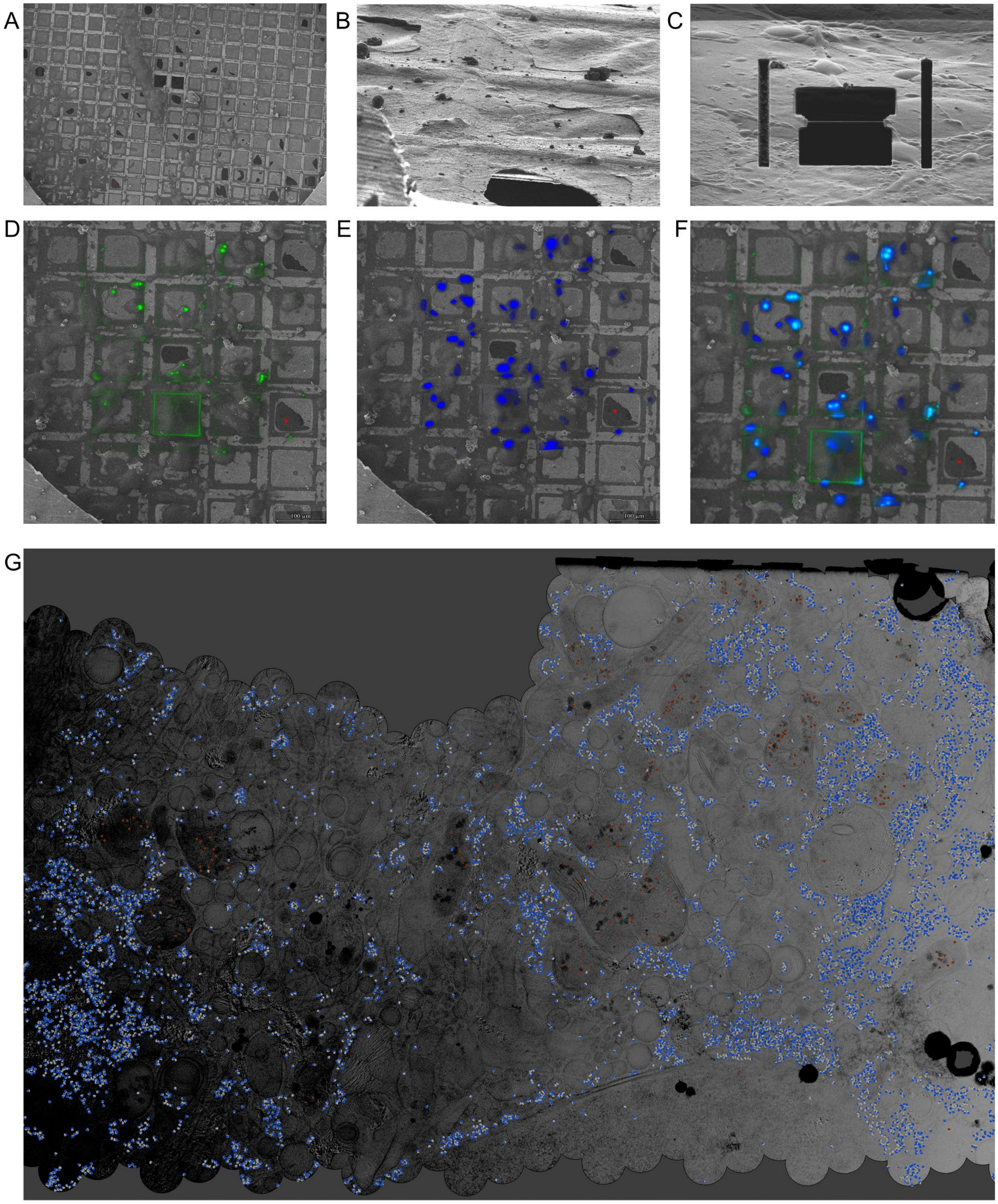

Figure S1

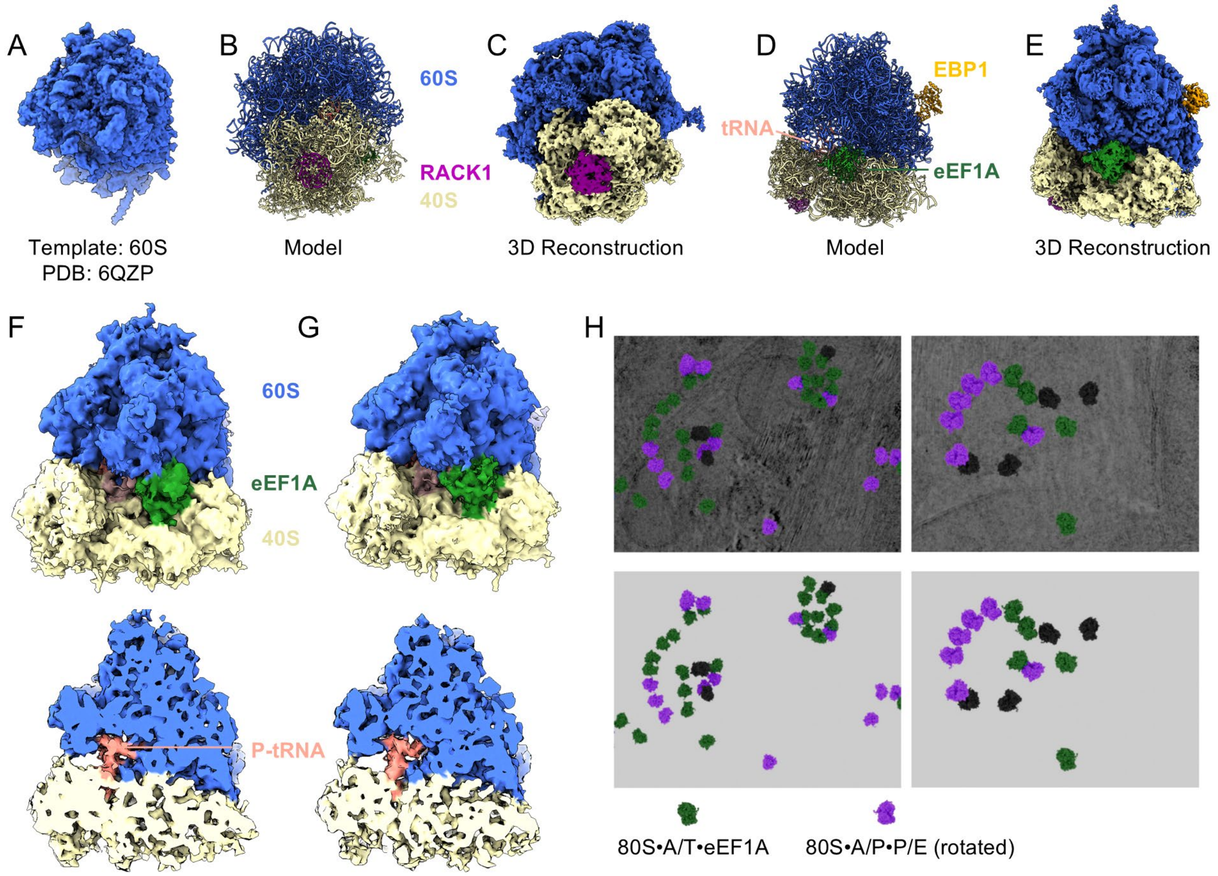

A
Template: 60S
PDB: 6QZP
B
60S
RACK1
40S
Model
C
3D Reconstruction
D
tRNA
EBP1
eEF1A
Model
E
3D Reconstruction
F
G
60S
eEF1A
40S
P-tRNA
H
80S•A/T•eEF1A
80S•A/P•P/E (rotated)

Figure S2

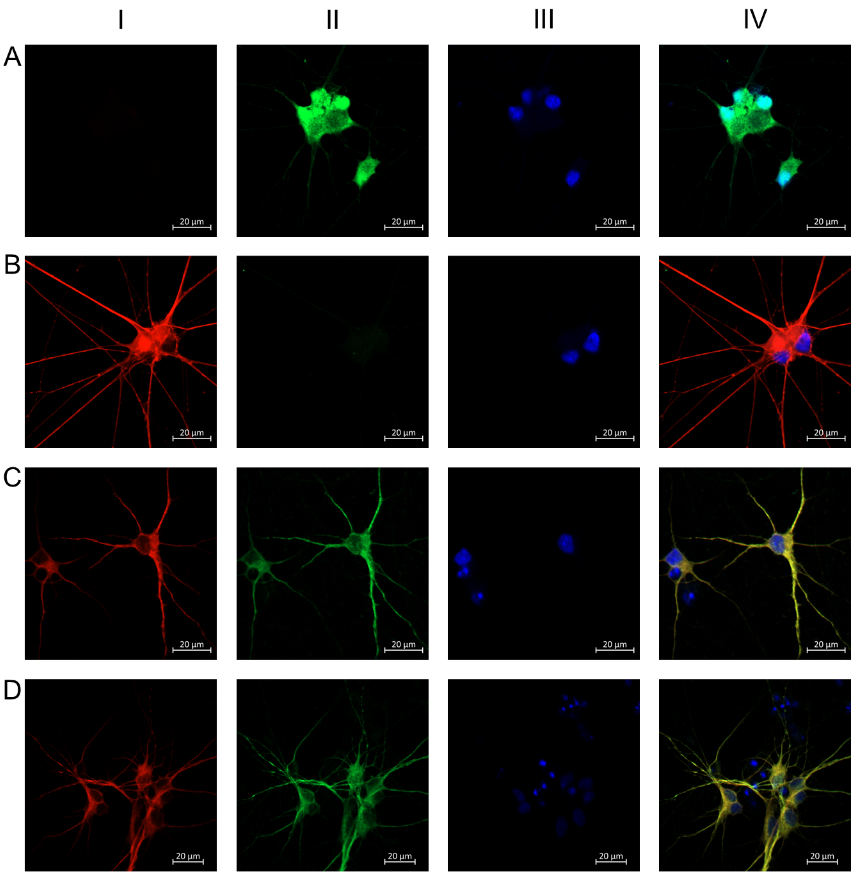

Figure S3

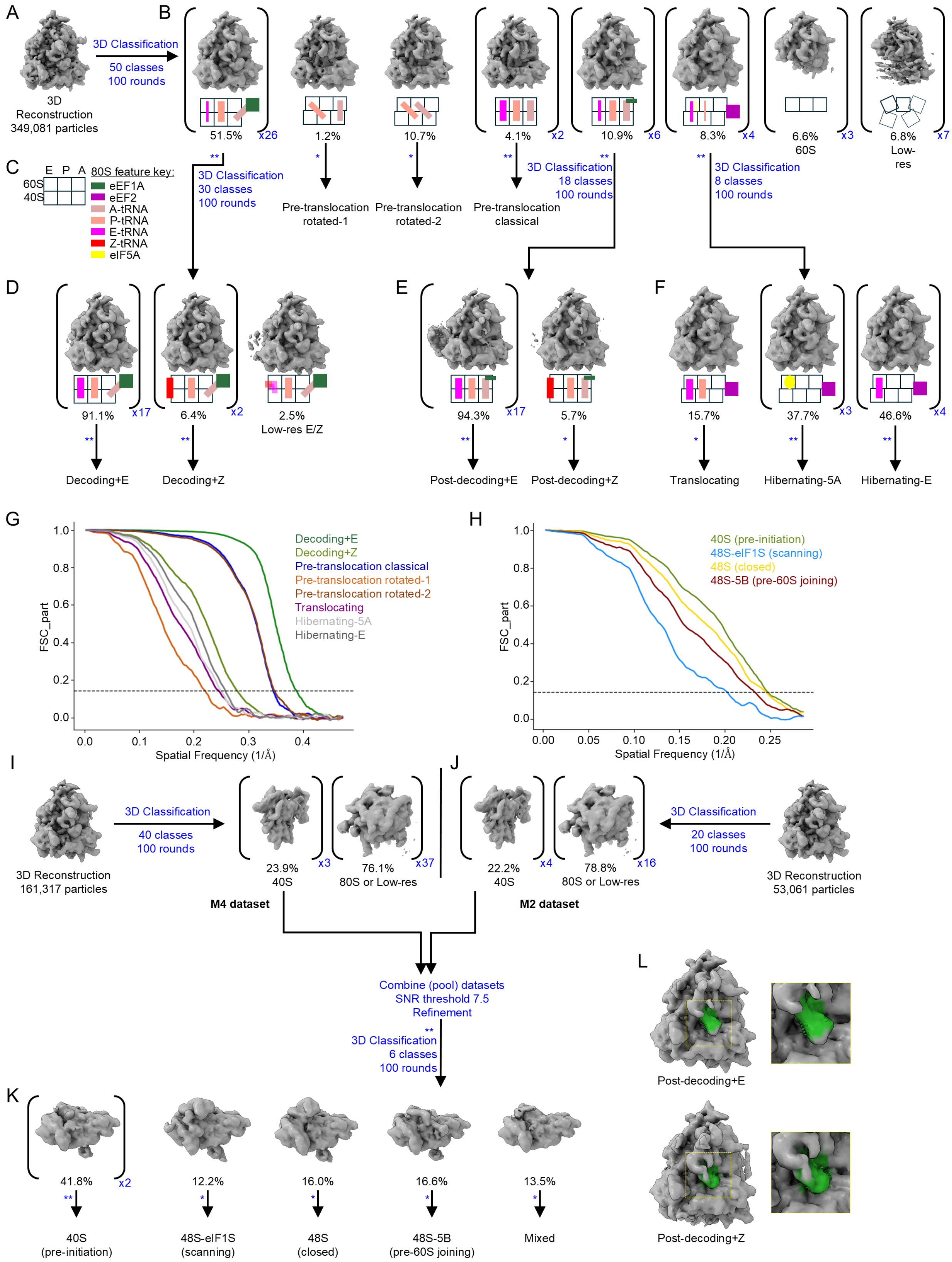

A
B
C
D
E
F
G
H
I
J
K
L
3D Reconstruction 349,081 particles
3D Classification
50 classes
100 rounds
51.5%
x26
1.2%
10.7%
4.1%
x2
10.9%
x6
8.3%
x4
6.6% 60S
x3
6.8% Low-res
x7
E P A 80S feature key:
60S
40S
eEF1A
eEF2
A-tRNA
P-tRNA
E-tRNA
Z-tRNA
eIF5A
3D Classification 30 classes 100 rounds
Pre-translocation rotated-1
Pre-translocation rotated-2
Pre-translocation classical
3D Classification 18 classes 100 rounds
3D Classification 8 classes 100 rounds
91.1%
x17
6.4%
x2
2.5% Low-res E/Z
94.3%
x17
5.7%
15.7%
37.7%
x3
46.6%
x4
Decoding+E
Decoding+Z
Post-decoding+E
Post-decoding+Z
Translocating
Hibernating-5A
Hibernating-E
Decoding+E
Decoding+Z
Pre-translocation classical
Pre-translocation rotated-1
Pre-translocation rotated-2
Translocating
Hibernating-5A
Hibernating-E
FSC_part
Spatial Frequency (1/Å)
40S (pre-initiation)
48S-eIF1S (scanning)
48S (closed)
48S-5B (pre-60S joining)
3D Reconstruction 161,317 particles
3D Classification
40 classes
100 rounds
23.9% 40S
x3
76.1% 80S or Low-res
x37
M4 dataset
22.2% 40S
x4
78.8% 80S or Low-res
x16
M2 dataset
3D Classification
20 classes
100 rounds
3D Reconstruction 53,061 particles
Combine (pool) datasets
SNR threshold 7.5
Refinement
3D Classification
6 classes
100 rounds
41.8%
x2
12.2%
16.0%
16.6%
13.5%
40S (pre-initiation)
48S-eIF1S (scanning)
48S (closed)
48S-5B (pre-60S joining)
Mixed
Post-decoding+E
Post-decoding+Z

### Figure S4

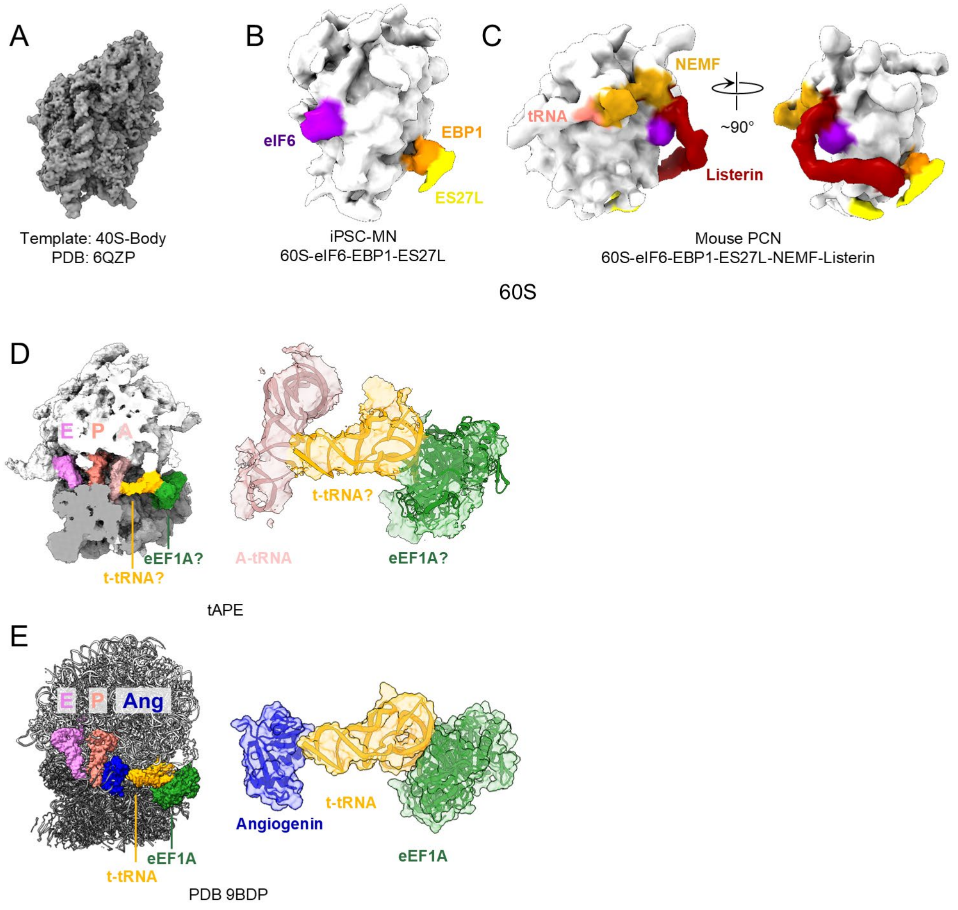

A
Template: 40S-Body
PDB: 6QZP
B
eIF6
EBP1
ES27L
iPSC-MN
60S-eIF6-EBP1-ES27L
C
NEMF
tRNA
~90°
Listerin
Mouse PCN
60S-eIF6-EBP1-ES27L-NEMF-Listerin
60S
D
E
P
A
eEF1A?
t-tRNA?
A-tRNA
t-tRNA?
eEF1A?
tAPE
E
E
P
Ang
eEF1A
t-tRNA
Angiogenin
t-tRNA
eEF1A
PDB 9BDP

Figure S5

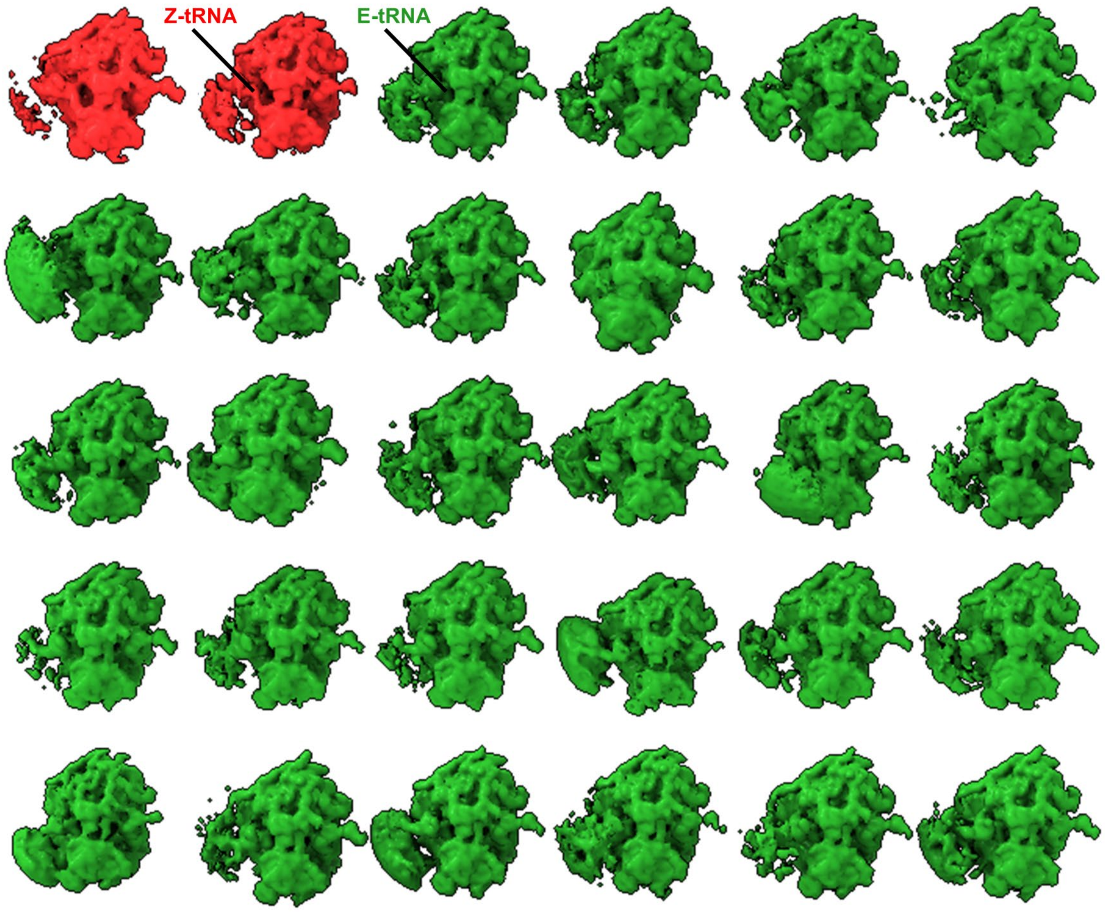

Figure S6

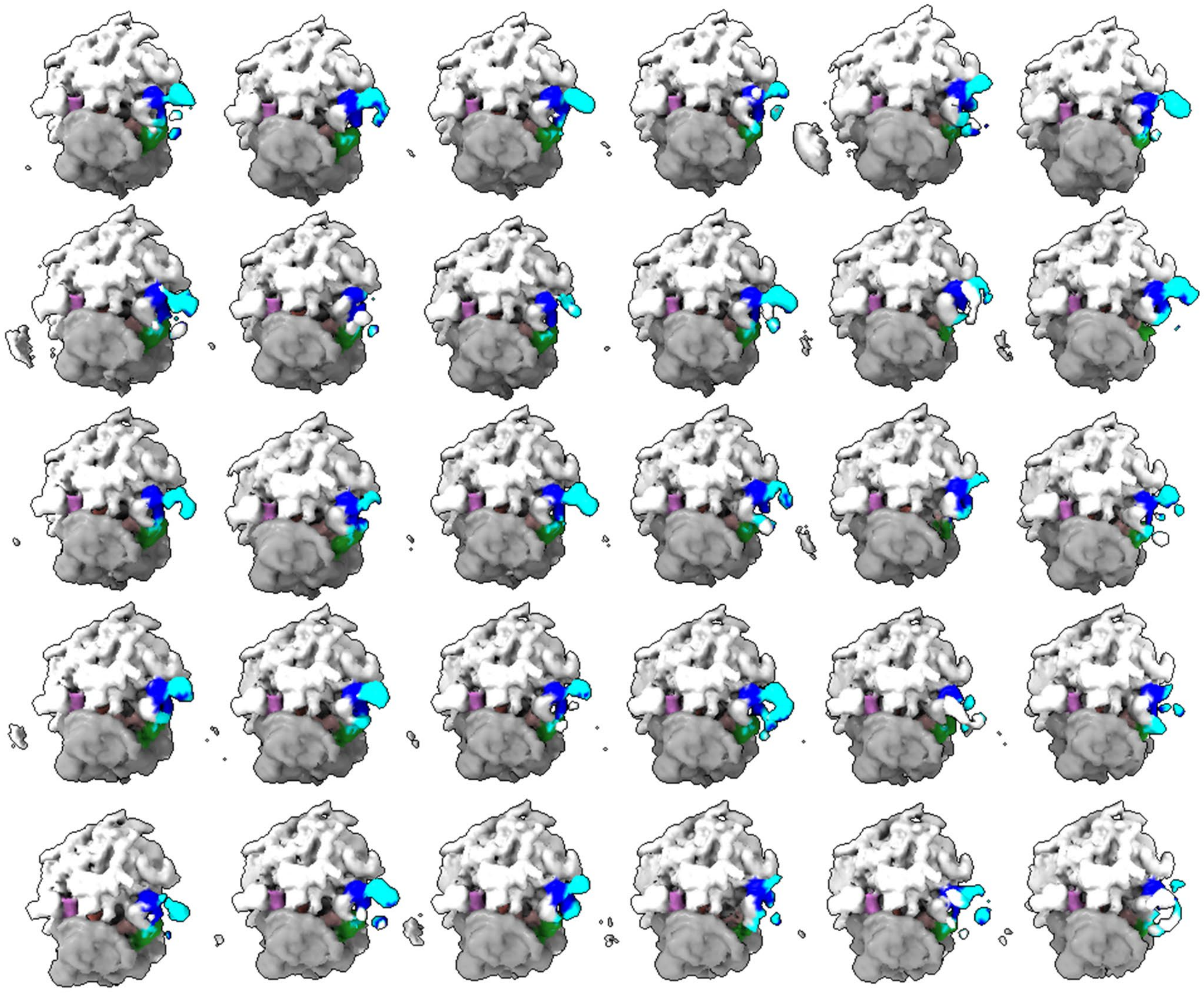

Figure S7

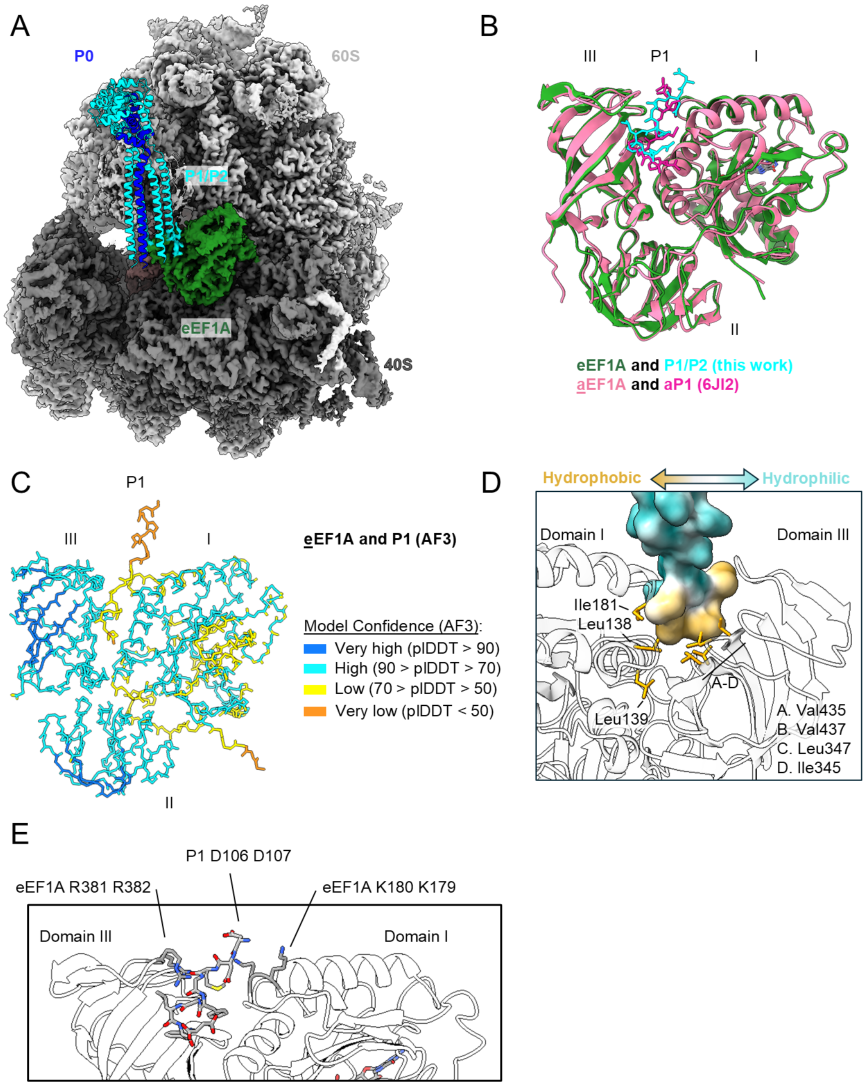

A
P0
60S
P1/P2
eEF1A
40S
B
III
P1
I
II
eEF1A and P1/P2 (this work)
aEF1A and aP1 (6JI2)
C
P1
III
I
II
eEF1A and P1 (AF3)
Model Confidence (AF3):
Very high (pIDDT > 90)
High (90 > pIDDT > 70)
Low (70 > pIDDT > 50)
Very low (pIDDT < 50)
D
Hydrophobic
Hydrophilic
Domain I
Domain III
Ile181
Leu138
Leu139
A-D
A. Val435
B. Val437
C. Leu347
D. Ile345
E
P1 D106 D107
eEF1A R381 R382
eEF1A K180 K179
Domain III
Domain I

Figure S8

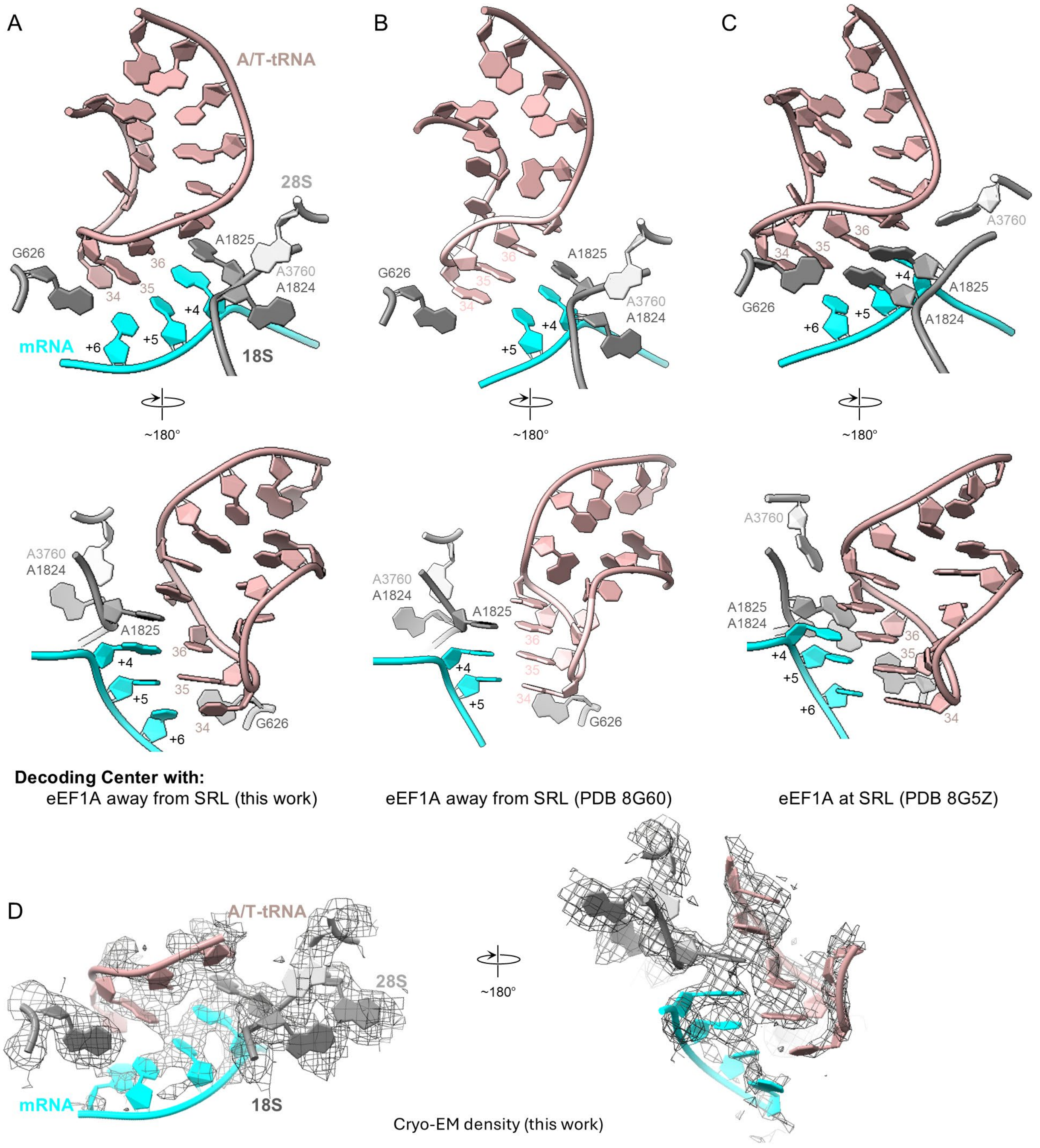

A
B
C
D
A/T-tRNA
28S
18S
mRNA
G626
A1825
A1824
A3760
34
35
36
+4
+5
+6
~180°
Decoding Center with:
eEF1A away from SRL (this work)
eEF1A away from SRL (PDB 8G60)
eEF1A at SRL (PDB 8G5Z)
Cryo-EM density (this work)

Figure S9

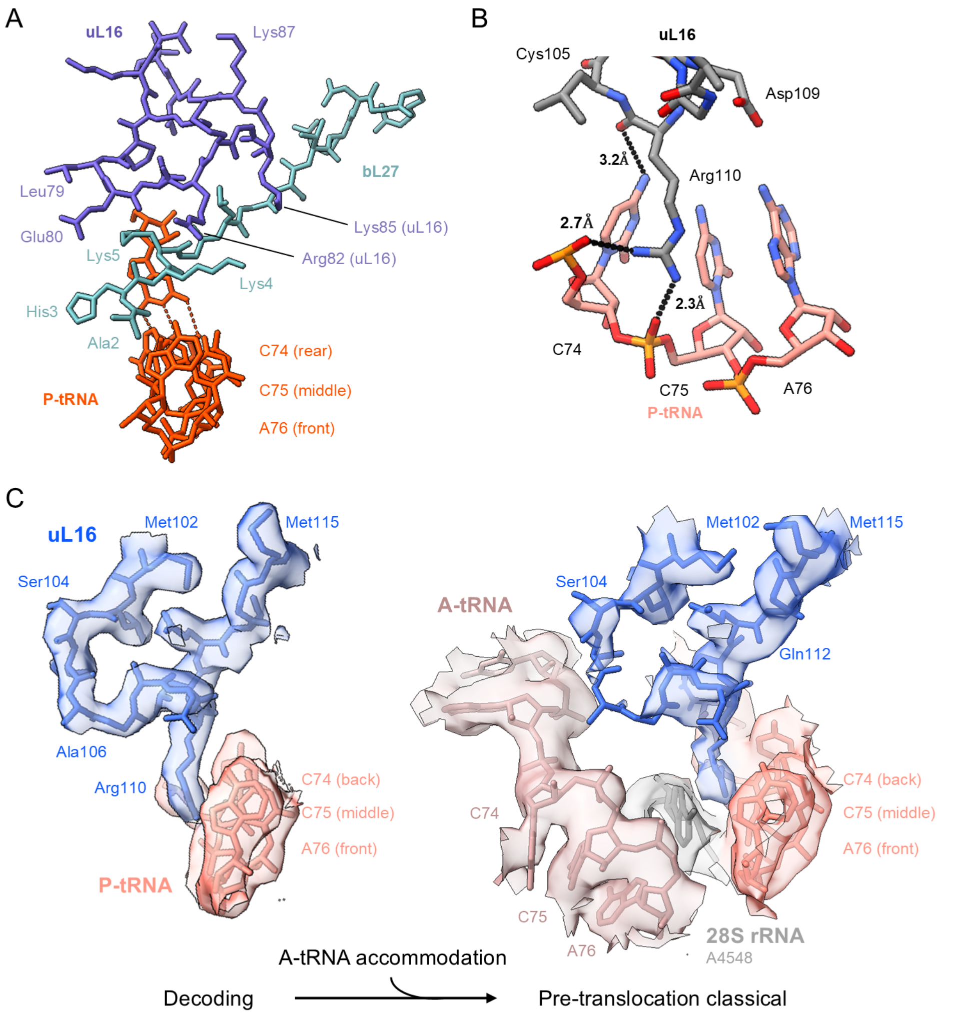

D

```
Archaea        MPLRPARCYTHFSGPPYTRREYIHGVPPPKITRFEMGNRE-MMEKWEIKGELVMLEAGQI 59
Yeast          MARRPARCYRYQKNKPYPKSRYNRAVPDSKIRIYDLGKKKATVDEFPLCVHLVSNELEQL 60
Human          MGRRPARCYRYCKNKPYPKSRFCRGVPDAKIRIFDLGRKKAKVDEFPLGGHMVSDEYEQL 60
Mouse          MGRRPARCYRYCKNKPYPKSRFCRGVPDAKIRIFDLGRKKAKVDEFPLCGHMVSDEYEQL 60
               *  ****** : .. ** : .: :.**  **  :::*.::  :::: :  .:*  *  *:

Archaea        RHNALEAARIAVNKYLATTIGDKNFYFRIRVYPHHVIRENKMMAFAGADRLQDGMRQAFG 119
Yeast          SSEALEAARICANKYMTTVSGRDAFHLRVRVHPFHVLRINKMLSCAGADRLQQGMRGAWG 120
Human          SSEALEAARICANKYMVKSCGRDGFHMRVRLHPFHVIRINKMLSCAGADRLQTGMRGAFG 120
Mouse          SSEALEAARICANKYMVKSCGKDGFHIRVRLHPFHVIRINKMLSCAGADRLQTGMRGAFG 120
                 :*******..***:..   *  . *::*:*::*.**:* ***:: ******* *** *:*

Archaea        KPVGTAARVYPGTVVLEVWARKGDEENVKEALRRGASKFPLPSRIVIRTRGAQAS----- 174
Yeast          KPHGLAARVDIGQIIFSVRTKDSNKDVVVEGLRRARYKFPGQQKIILSKKWGFTNLDRPE 180
Human          KPQGTVARVHIGQVIMSIRTKLQNEEHVIEALRRAKFKFPGRQKIHISKKWGFTKFNADE 180
Mouse          KPQGTVARVHIGQVIMSIRTKLQNKEHVIEALRRAKFKFPGRQKIHISKKWGFTKFNADE 180
               ** *  .***   * :::.: ::   ::: * *.***.   ***   .:* : .: . :.

Archaea        ------------------------------------------      174
Yeast          YLKKREAGEVKDDGAFVKFLSKKGSLENNIREFPEYFAAQA       221
Human          FEDMVAKKCLIPDGCGVKYVPSHGPLDKWRVLHS-------       214
Mouse          FEDKVAAKRLIPDGCGVKYIPERGPLDKWRALHS-------       214
```

Figure S10

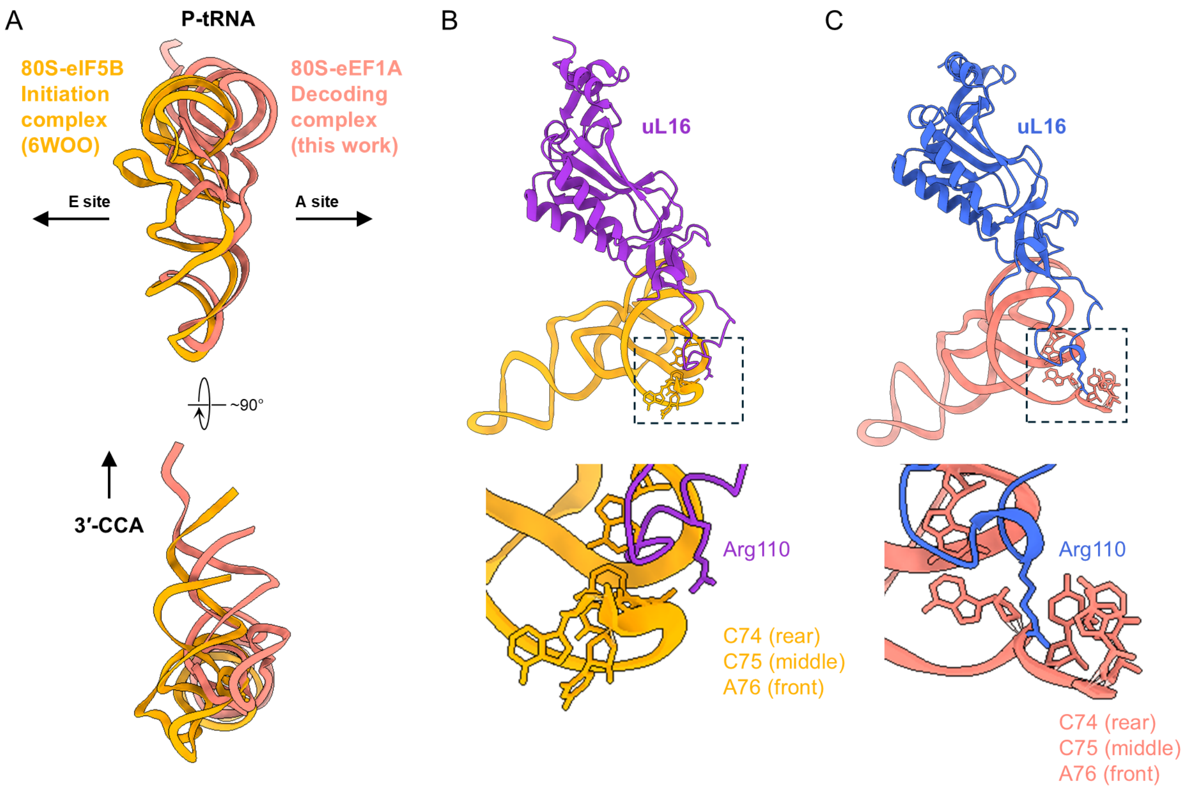
A
P-tRNA
80S-eIF5B Initiation complex (6WOO)
80S-eEF1A Decoding complex (this work)
E site
A site
~90°
3′-CCA
B
uL16
Arg110
C74 (rear)
C75 (middle)
A76 (front)
C
uL16
Arg110
C74 (rear)
C75 (middle)
A76 (front)

Figure S11

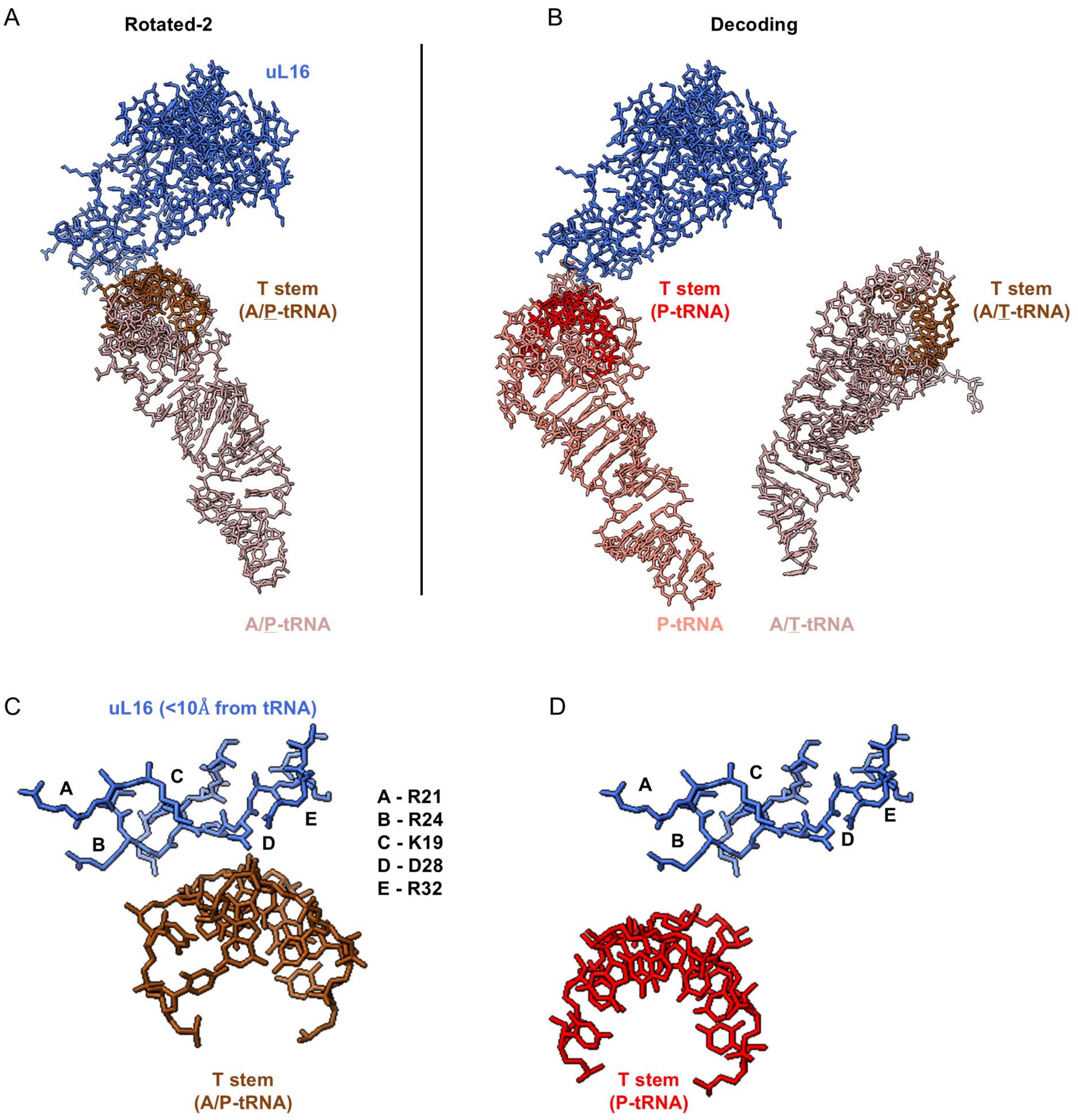
A
Rotated-2
uL16
T stem
(A/P-tRNA)
A/P-tRNA
B
Decoding
T stem
(P-tRNA)
T stem
(A/T-tRNA)
P-tRNA
A/T-tRNA
C
uL16 (<10Å from tRNA)
A
B
C
D
E
A - R21
B - R24
C - K19
D - D28
E - R32
T stem
(A/P-tRNA)
D
A
B
C
D
E
T stem
(P-tRNA)

Figure S12

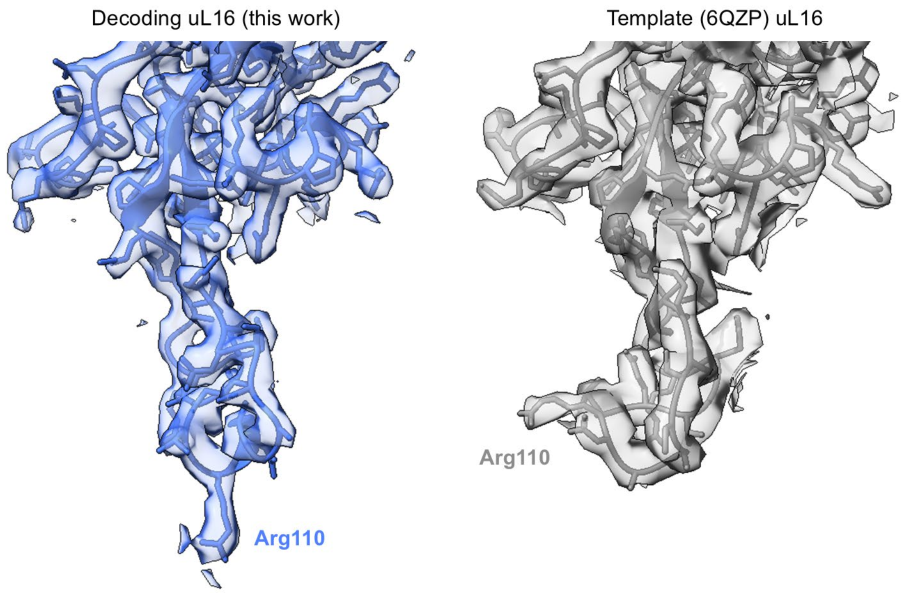

Figure S13

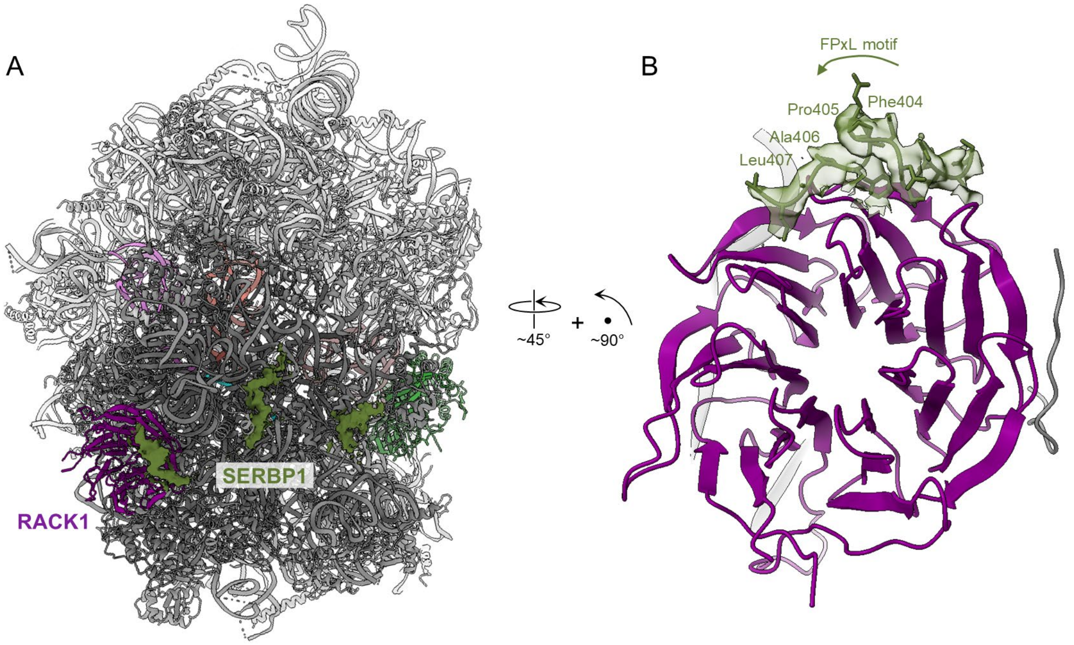

A
SERBP1
RACK1
~45°
~90°
B
FPxL motif
Pro405
Phe404
Ala406
Leu407

Figure S14
